# Adversarial learning enables unbiased organism-wide cross-species alignment of single-cell RNA data at scale

**DOI:** 10.1101/2024.08.11.607498

**Authors:** Juan Javier Díaz-Mejía, Elias Williams, Brendan Innes, Octavian Focsa, Dylan Mendonca, Swechha Singh, Allison Nixon, Ronen Schuster, Matthew B. Buechler, Boris Hinz, Sam Cooper

## Abstract

Today’s single-cell RNA (scRNA) datasets remain siloed, due to significant challenges associated with their integration at scale. Moreover, most scRNA analysis tools that operate at scale leverage supervised techniques that are insufficient for cell-type identification and discovery. Here, we demonstrate that the alignment of scRNA data using unsupervised models is accurate at an organism-wide scale and between species. To do this, we show adversarial training of a deep-learning model we term batch-adversarial single-cell variational inference (BA-scVI) can be employed to align standardized benchmark datasets comprising dozens of scRNA studies spanning tissues in humans and mice. In the aligned space, we analyze cell types that span tissues in both species and find prevalent complement expressing macrophages and fibroblasts. We provide access to the tools presented via an online interface for atlas exploration and reference-based drag-and-drop alignment of new data.

## Introduction

Single-cell RNA sequencing (scRNA) is able to dissect tissue and experimental models with unprecedented precision and is underpinning a wave of new biological discovery. Today, scRNA analysis focuses on individual datasets or a handful of datasets with similar etiology. Yet, to build a comprehensive organism-wide understanding of gene expression profiles underlying cell types and stages we will need to examine integrated transcriptional atlases that combine studies and patient populations at scale (Regev et al. 2017). The ever-growing number of published scRNA studies creates an opportunity for the development of a large aligned scRNA atlas (Gavish et al. 2023), that would enable standardized reference based analysis, and seamless cross-dataset comparison (Lotfollahi et al. 2024). However, the challenge of combining data from multiple disparate scRNA studies remains (Butler et al. 2018; Gavish et al. 2023; Lotfollahi et al. 2024; Lähnemann et al. 2020).

While studies have looked at the alignment of batches within individual datasets or a collection of integration tasks, none have focused on how well models align studies at scale, especially across tissue types, instruments, and species as would be required for generation of a reference atlas (Huang et al. 2021; Xie et al. 2021; Abdelaal et al. 2019; Diaz-Mejia et al. 2019; Christensen et al. 2023; Butler et al. 2018; Song et al. 2023). Moreover, those studies that have used models across tissue types or studies have used supervised models trained on cell-type labels, such as scBERT, Celltypist, and SCimilarity (Yang et al. 2022; Domínguez Conde et al. 2021). Yet, unsupervised alignment is superior for cell-type discovery (Vasighizaker, Danda, and Rueda 2022) and will be needed for unbiased cross-species comparative analysis. Thus, a significant demand exists for a reference atlas and reference analysis based on unsupervised alignment (Lotfollahi et al. 2024). In this study, we leverage the analysis of a large human scRNA benchmark dataset to test the ability of methods to align scRNA data between studies and tissue types first and then between species using a mouse atlas. In showing that top models can accurately align the atlases with minimal loss to cell-type granularity, we demonstrate that reference-based analysis is possible with a single unsupervised model and that cell types can be compared across tissues and between species, paving the way to phylogenetic cell-type analyses. Finally, we provide this model, an online tool for exploring the results, and a tool for drag-and-drop alignment of new data to give the broader community access to the work we present here.

## Results

### Construction and state-of-the art alignment of the scREF scRNA benchmark dataset

In scRNA studies, unsupervised models transform a high-dimensional gene-expression space into a low-dimensional cell-type space. We sought an optimum model for performing this task while aligning scRNA datasets across publications. To test alignment performance, we thus developed the scREF benchmark, a collection of 46 human scRNA studies, spanning 2,359 samples and 36 tissues, where for each dataset, quality checks have been performed and metadata standardized (Figure 1a; Methods). In scREF, we include organ-specific and human-wide datasets, e.g., the Tabula Sapiens (Tabula Sapiens Consortium* et al. 2022) and the Human Cell Landscape (Han et al. 2020). Importantly, scREF includes data from droplet-based (10X 5’, 10X 3’, 10X multiome, and Dropseq) and plate/bead based methods (Microwell-seq, Seq-Well and SMARTScribe) to test cross-technology alignment. Author-provided cell-type labels for 45 studies were acquired and standardized, while for three cases, we generated labels reproducing the original author’s pipeline (Table S1; Methods); overall, this resulted in 60 unique cell-type labels. Tissue-type labels were standardized for plotting and analysis (Table S1). For training and testing, we balanced dataset representation by stratifying cell types from each tissue type in each dataset (Table S1; Methods), leading to the final 1.21 million cell evaluation benchmark.

**Figure 1.**
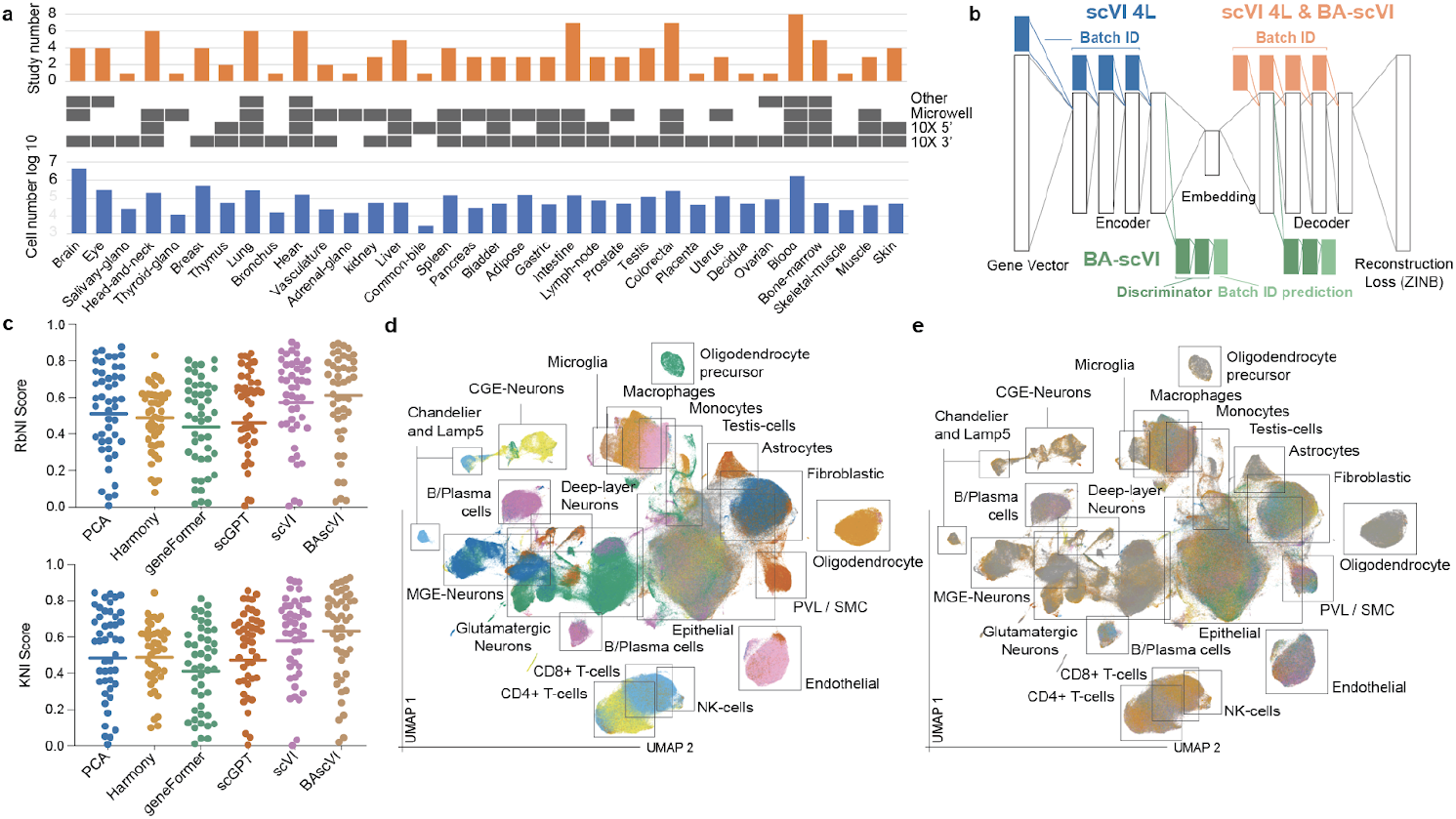
Alignment of a human scRNA reference atlas: a) Summary statistics of the scREF dataset, broken down by tissue type and instrument; b) Architecture of the optimized scVI and BA-scVI models; c) KNI and RbNI scores were determined for the aligned scREF dataset, based on standardized author cell type labels for the alignment tools. Data points correspond to the average score achieved by the model on a study. The average score obtained on the entire benchmark plotted as a line; d) A UMAP projection of the aligned stratified scREF atlas (n=1.27m), coloured by ‘ground-truth’ standardized author cell-type label. The legend is omitted for brevity (coloring is the same as Figure 3d), boxes show major cell-type groupings; e) same projection as (d), coloured by study name the legend is omitted.

We aimed to identify models that most effectively remove technical batch effects while aligning cell types. We thus sought a metric to compare model performance for this task. We developed two metrics that we term K- and Radius-based Neighbors Intersection scores (KNI and RbNI; Methods; Supplementary) that combine the kBET score for batch-effect detection (Büttner et al. 2019) with cross-dataset cell-type prediction accuracy of author labels, a gold-standard metric for preservation of biological signal (Domínguez Conde et al. 2021). In developing the KNI and RbNI scores, we evaluated these and other benchmark metrics on simulated data, real data with synthetic batch effects or noise, and in a real-world setting on a small scRNA benchmark that we also used for model optimization/ development (Supplementary; Table S2). Across these analyses, we find that the KNI and RbNI metrics capture the quality of cell-type space in a single value, providing a simple, robust performance readout.

Following initial optimization, we tested the ability of published, scalable scRNA analysis models to align the scREF atlas. Here we found that an optimized variant of single-cell Variational Inference (scVI)(Lopez et al. 2018) outperformed Harmony (Korsunsky et al. 2019), PCA on highly variable genes, geneFormer fine-tuned for batch effect correction with scVI (Theodoris et al. 2023) and scGPT fine tuned for batch effect correction as per the author protocol (Cui et al. 2024) (Figure 1c; (Lopez et al. 2018)). Qualitatively, UMAP projections showed that scVI produces a reasonably high degree of alignment accuracy (Figure S8). Significantly, organism-wide studies from markedly different technologies Microwell-seq (Han et al. 2020), and 10X (Tabula Sapiens Consortium* et al. 2022) overlap extensively with each other and have KNI/RbNI scores at or above average (Table S3), indicating alignment independent of technology. We thus find that an optimized scVI model can be used to perform effective large-scale alignment. However, we noted that the direct penalization of batch effects performed in Harmony can improve batch effect correction (Supplemental).

Adversarial learning has emerged as a powerful approach to identifying and removing technical artifacts in generative machine-learning outputs (Goodfellow et al. 2020). We developed batch adversarial scVI (BA-scVI) as a variant of scVI, leveraging an adversarial training approach to remove technical artifacts (i.e., batch effects) in scRNA data at train time, similar to that used for the cell-search algorithm in cellBLAST (Cao et al. 2020; Shaham 2018). In BA-scVI, a discriminator learns to predict the batch identifier from the encoder output and decoder input in the first update step (Figure 1b). In the second update step, a modified scVI architecture seeks to minimize reconstruction loss and KL-divergence while maximizing discriminator loss, where discriminator loss represents the discriminator’s ability to predict the batch identifier (Figure 1b; Methods). By maximizing this discriminator loss while minimizing reconstruction loss, the model directly penalizes batch effects while rewarding preserving cell-type information in the latent space. In line with this, an optimized BA-scVI model outperforms scVI on scREF (Figure 1c). This leads to effective cross study alignment, clearly resolved cell-type clusters in qualitative UMAP projections of the embedding space (Figure 1d, e).

### Alignment of the scREF benchmark dataset maintains cell-type granularity

A major concern in the atlas-building community is that aligning datasets reduces the granularity of cell-type detection. To qualitatively assess how well cell-type labelings are preserved at the organ level in the aligned cell-type space, we fit UMAP to the three best-represented tissues: breast (4 studies), brain (4 studies), and blood (7 studies). Supporting effective alignment with BA-scVI, we found effective distinction of standardized cell types indicating preserved granularity (Figure 2a), alongside significant overlap between studies (Figure 2b). BA-scVI could also resolve ‘original author’ labels in UMAP projections of an example study for each tissue type, qualitatively supporting the preservation of cell-type resolution (Figure 2c) in the aligned space.

**Figure 2.**
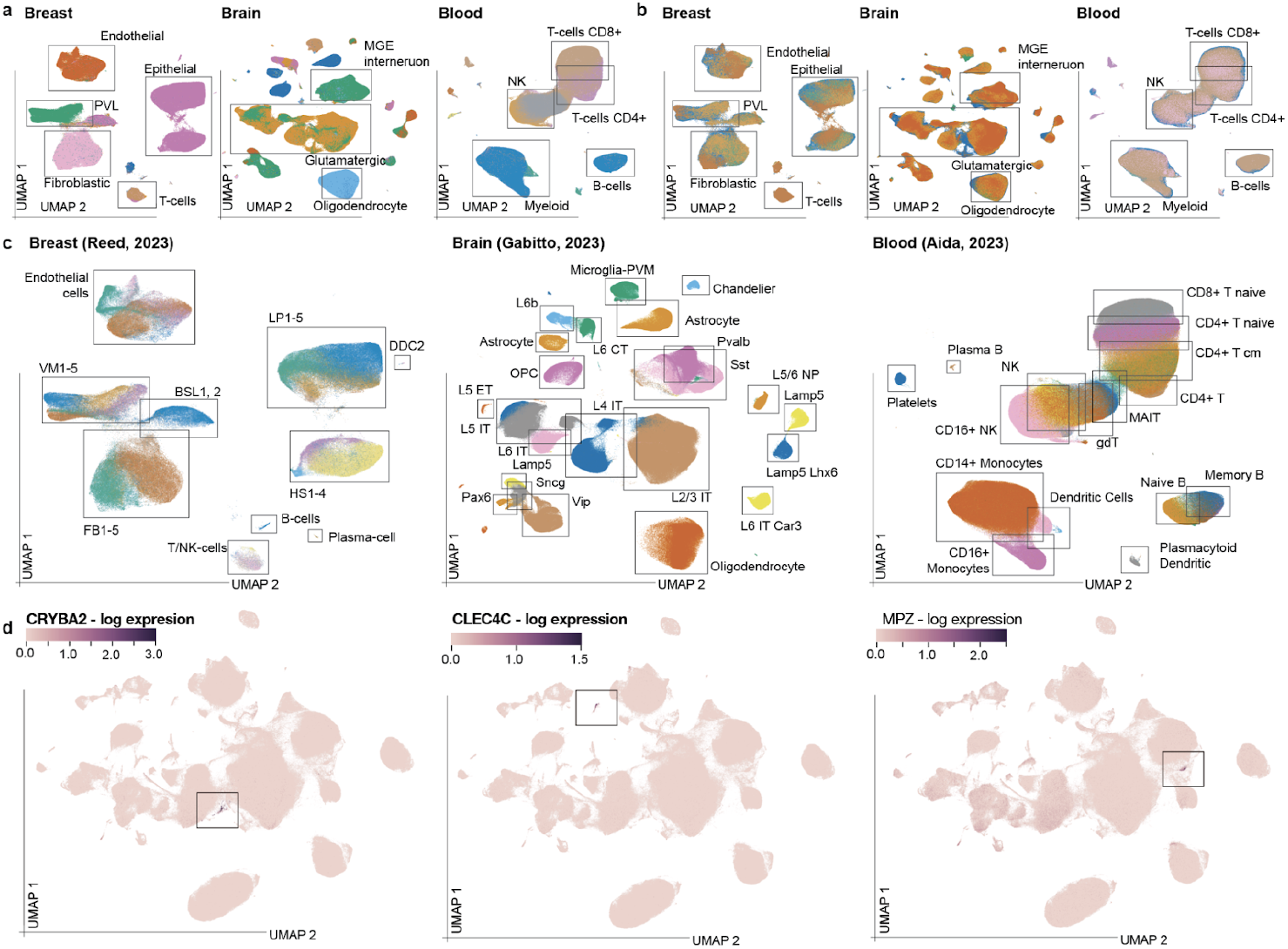
BA-scVI scREF maintains cell-type granularity on alignment: a) 10-dimensional scRNA embeddings from BA-scVI corresponding to Breast (n=0.4m cells), Brain (n=4.8m cells), and Blood (n=1.6m cells) tissue-types were projected into a 2-dimensional space with UMAP. Cells are coloured by the standardized ground-truth labels, highlighting the consistency with which BA-scVI is able to align similar cell types. The cell-type and study legends here and in (b/c) are omitted for brevity; major groupings are in boxes; b) The same UMAP projections coloured by study name; c) The same UMAP projections coloured by original author labels for example studies; Breast (n=0.3m cells), Brain (n=0.8m cells), Blood (n=1m cells); d) scREF UMAP projections coloured by expression of CRYBA2, CLEC4C, and MPZ; selective markers of colorectal endocrine, plasmacytoid dendritic, and Schwann cells respectively.

Quantitatively, using a KNN accuracy test with 2-fold cross-validation, for example, studies (Methods), we found that relative cell-type embedding localizations of the original author labels are conserved since the model can obtain high cell-type labeling accuracies on held-out data. Specifically, KNN accuracies of; (1) 83% were obtained on a large breast dataset (Reed et al. 2023); increasing to 96% on ‘numerical’ subtype merging (e.g. cell subtypes ‘LP1’ to ‘LP5’ become ‘LP’); 2) 99 % for a brain study Gabitto et al. (Gabitto et al. 2023); and 3) 83% for the Kock et al. blood dataset where T-cell subtype label overlap is notably seen in projections in the original study (Kock et al. 2024). Overall this analysis further supports preservation of cell-type granularity. Furthermore, we also find that rare cell types can be distinguished in BA-scVI alignments. Specifically, colorectal, endocrine cells, plasmacytoid dendritic cells, and Schwann cells are marked by CRYBA2, CLEC4C, and MPZ, respectively (Supplemental). In the aligned atlas, selective groupings of these genes can be seen, indicating these rare cell-type groupings are captured effectively (Figure 2d). Finally, we see that BA-scVI performs well at the identification of cell subtypes in a discovery setting (Supplemental). Overall, these results provide strong evidence that cell-type granularity is preserved following alignment with BA-scVI, supporting its power as a model for unsupervised reference-based scRNA analysis.

### Cross-species alignment of scRNA reference atlases can improve accuracy in data-poor species

Next, we tested whether we could use BA-scVI to align atlases between species. For this, we constructed scREF-mu, comprising 18 mouse scRNA studies, 1,290 samples, 34 tissues, and over 3 million cells, with author cell-type labels covering 11 studies (Table S1). As in scREF, each tissue type included was required to appear in at least two datasets. However, we noted that in contrast to the scREF human dataset, different mouse brain-tissue regions were often only represented abundantly in one dataset, e.g., the hippocampus (Yao et al. 2021), and cerebellum (Kozareva et al. 2021). We trained BA-scVI on scREF-mu alone and jointly with the scREF human dataset. KNI and RbNI scores were calculated for scREF-mu alone, jointly trained using mouse cell-type labels, and jointly trained using both human and mouse labels. Here, we found that joint training with the inclusion of cell-type labels from both species resulted in the highest KNI and RbNI scores (Figure 3a; Table S4), indicating value in cross-species alignment. Notably, the KNI and RbNI scores on the human scRNA datasets did not decrease (Figure 3b), and qualitative assessment of UMAP projections of the aligned atlas also supports successful alignment (Figure 3c, d). Cerebellar granule cells only appear once in both the mouse and human atlases and are thus excluded in the benchmarks. Yet, we find we can accurately label these cells leveraging cross-species alignment (Supplemental; Figure S11). Overall, we show that cross-species alignment of organism-wide scRNA atlases is possible and may improve the labeling of cell types in data-poor species.

**Figure 3.**
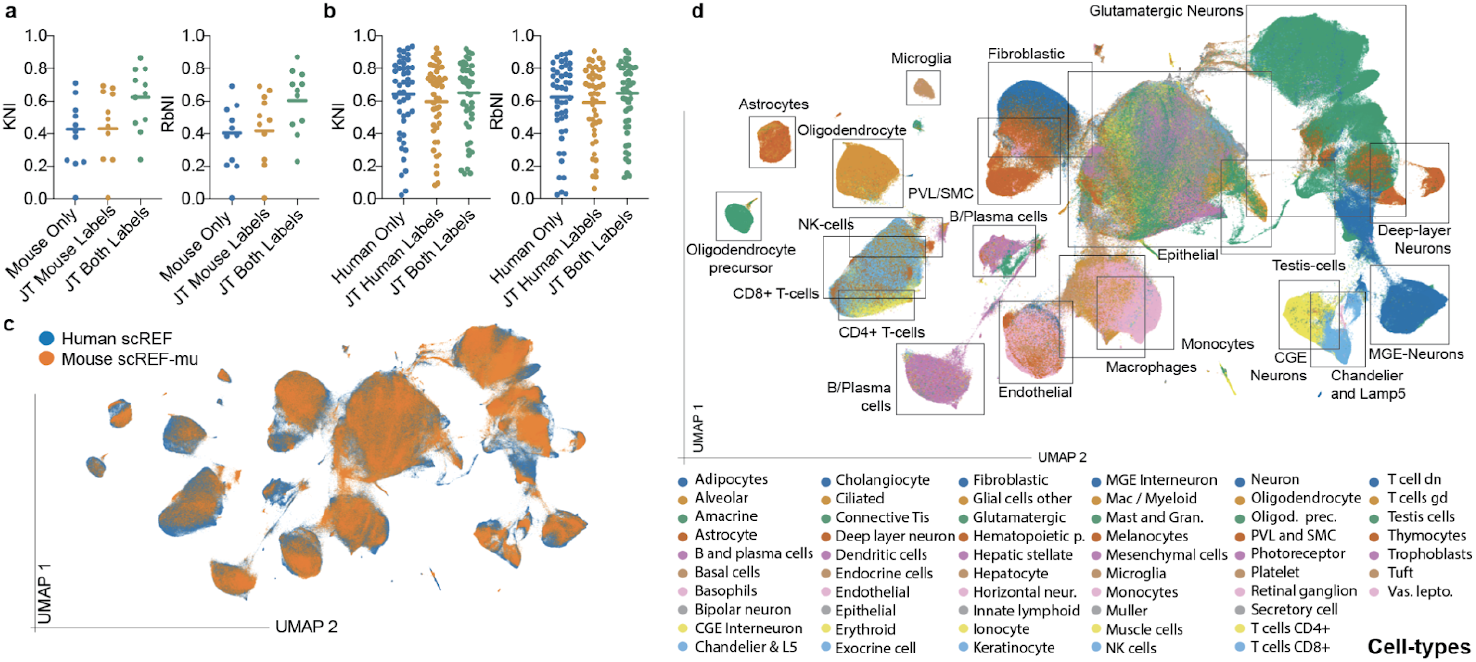
Cross-species alignment of mouse and human scRNA atlases: a) Analysis of KNI and RbNI scores obtained by BA-scVI on alignment of the mouse scRNA atlas scREF-mu only (Mouse Only), jointly trained with assessment of alignment accuracy using mouse labels only (JT Mouse Labels), and jointly trained leveraging labels from both mouse and humans for accuracy assessment (JT Both Labels); b) KNI and RbNI scores from BA-scVI alignment of the human scRNA atlas scREF only (Human Only) and jointly trained with assessment of alignment accuracy using human labels only (JT Human Labels), and jointly trained leveraging labels from both mouse and humans for accuracy assessment (JT Both Labels); c) UMAP projections of the jointly aligned atlas, coloured by organism with Mouse (Orange; n=528k), humans (Blue; n=1.27m); d) UMAP projection of the jointly aligned atlas coloured by cell-type, with major groupings highlighted by boxes.

### Cross-species alignments enables us to analyze cell-type conservation across tissues and species

Following organism-wide alignment, we looked to understand how cell types group together in the aligned atlas using an unbiased approach. We tested both K-means clustering and Label Propagation (LP) approaches ability to identify cell-type groupings on the aligned scREF and scREF-mu datasets (∼14m cells) as unbiased methods that could scale millions of cells (Methods). We found that LP results aligned well with the original author as well as standardized labels, compared to K-means, as measured by the Adjusted Rand Index (ARI), both on this dataset and the individual scREF and scREF-mu datasets (Figure 4a; Figure S12). Notably, we found the ARI between LP groupings and standardized labels was similar to that between standardized labels and author labels, giving us confidence this method of clustering generates meaningful groupings. Overall, LP identified 792 unique clusters in the cross-aligned atlas where more than 50 cells were present in at least two datasets. Approximately half of these clusters were CNS cell types (Figure 4b), which is in line with the enormous cell-type diversity seen in the brain (Zhang et al. 2023). The other half corresponded to the remaining tissue types, in line with prior estimates that hundreds of cell types exist in the body (Hatton et al. 2023). Similar results were obtained for the individual atlases (Supplemental).

**Figure 4.**
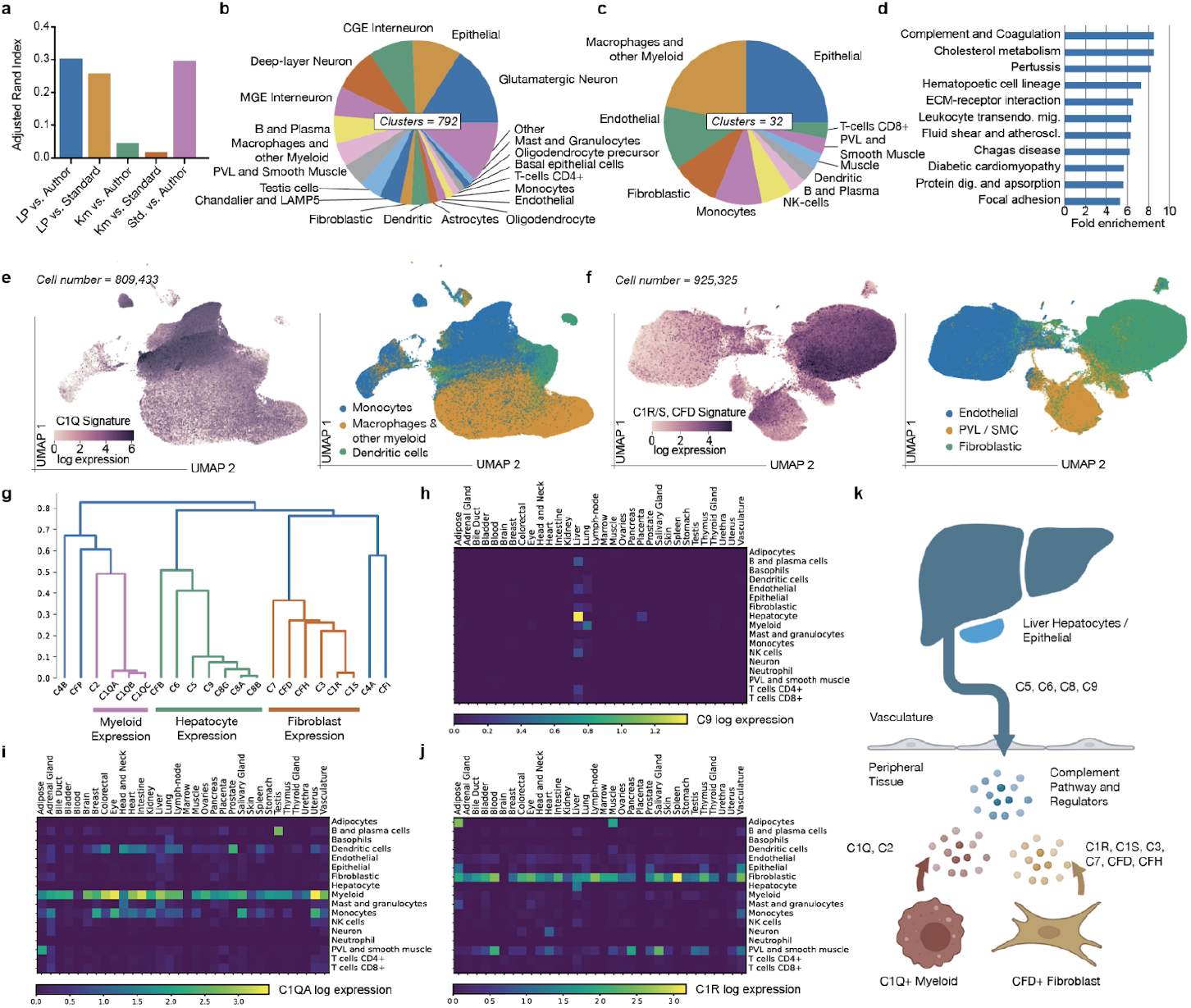
Analysis of conserved cell-types in the body: a) Comparison of label propagation and K-means clusters with author or standardized cell-type labels, measured by the Adjusted Rand Index for the jointly aligned mouse and human atlas; b) Proportion of unique cell-type groups, with more than 20 cells present in two datasets, by standardized label in the jointly aligned atlas; c) Same as (b) but filtered for cell-types with over 20 cells present in 2 datasets, in more than 10 tissues in mice and humans; d) Pathway enrichment for the top 20 differentially expressed genes in the conserved cell-types; e) UMAP projection of Myeloid lineages, coloured by mean C1QA, C1QB, and C1QC read count (left), or standardized cell-type label (right), n=695,141 cells; f) UMAP projection of Fibroblastic, Endothelial, and PVL / Smooth muscle cell types, colored by mean C1R, C1S, read counts (left), or standardized cell-type label (right); g) Hierarchical clustering of complement expression patterns h-j) Heatmaps of log read-counts of representative expression profiles for the three observed patterns; g) C9 = Liver; h) C1QA = Myeloid-types; i) C1R = Fibroblasts); k) Cartoon of proposed cell-type contributors to complement signaling.

Where most studies have focused on organ-specific cell-type ID, here we sought to characterize cell-type groupings that span tissue types and are highly conserved between organisms. We reason this may indicate evolutionary constraint. Clusters were filtered for those containing cells from 10 or more tissues in both the mouse and human datasets; potential artifactual groupings were also removed (Methods). After this, 32 conserved cross-tissue clusters remained, covering a diverse range of cell types, as marked by the modal standardized cell-type label (Figure 4c). We then leveraged differential gene-expression analysis to understand if these cell types are all well-defined or if new groupings emerge from this cross-tissue analysis (Methods). While most cell types were well defined, we noted macrophage groupings that show highly selective C1Q expression and fibroblast clusters that are enriched for C1R, C1S, and CFD, and SERPING1 (complement inhibitor) expression in the cross-aligned and individual atlases (Table S5). Pathway analysis (Ge, Jung, and Yao 2020) of the top 25 most-selective genes to each cell type in the cross-species returns “Complement and coagulation cascade” as the most enriched pathway in this gene set (Figure 4d), indicating this enrichment is not just qualitative.

While it is known that peripheral macrophages and dendritic cells express C1Q (Castellano et al. 2010; Müller, Hanauske-Abel, and Loos 1978), these genes are not typically used as myeloid cell-type markers. Moreover, while studies have shown that synovial fibroblasts can synthesize complement proteins in vitro (Katz and Strunk 1988), widespread fibroblast expression of complement has not been reported, although C7 is noted as a fibroblast markers (Buechler et al. 2021; Dominguez et al. 2020). Supporting the results obtained from label propagation, we found in UMAP projections of myeloid cell types, C1Q expression marks a distinct region (Figure 4d. Similarly, a C1R, C1S, and CFD signature marks an axis of variation in fibroblast cell types (Figure 4e). Expression of the core complement gene family in the unaligned scREF and scREF-mu atlas further supports our findings and also hints at broader complement regulation by fibroblasts and macrophages. Specifically, in the scREEF and scREF-mu atlas, we see three expression patterns emerge across standardized cell types and tissues (Figure 4f-i). Namely, 1) Myeloid selective expression of C1Q genes and C2; 2) Fibroblast selective expression of C1R, C1S, C3, CFH, CFD, and C7 with trace endothelial and PVL/SMC readouts; and 3) Liver / Hepatocyte selective expression of C5, C6, C8 components, C9 and CFB; with outlier expression of C4A, C4B, CFI and CFP. Importantly, these results are also seen in the individually aligned mouse and human atlases (Supplemental; Figure S13, 14). Overall, this analysis thus finds peripheral complement mRNA expression in broadly abundant, evolutionarily conserved fibroblasts and macrophages and highlights the value of the cross-species alignment we perform here in uncovering these expression patterns.

### BA-scVI trained on scREF realizes drag-and-drop scRNA analysis

Unlike scVI, BA-ScVI only uses batch labels in training. This allows new samples to be embedded de novo, as long as the contributing tissue is currently represented in the atlas. We demonstrate this at scale by using BA-scVI trained on scREF without the Tabula Sapiens atlas to accurately predict cell-type labels in the Tabula Sapiens atlas without further fine-tuning or training (Tabula Sapiens Consortium* et al. 2022)(Supplemental, Figure S15). We also found that this approach can map in vitro cell types to tissue scRNA data, which is valuable for in vitro model relevance optimization (Supplemental). To give broad access to this capability, we provide an online tool for drag-and-drop embedding and labeling of new human scRNA data against scREF. This tool is available as a user-friendly web app that also facilitates analysis and visualization at https://scref.phenomic.ai/. An API alongside R and Python functions is also given in the data availability section below.

## Discussion

Most scRNA data alignment benchmarking studies have used a handful of datasets to evaluate the performance of machine-learning methods for this task (Pasquini et al. 2021; Abdelaal et al. 2019; Diaz-Mejia et al. 2019). However, the development of reference-based atlases will require the alignment of hundreds to thousands of samples and millions of cells to one another. In this study, we present scREF as a large-scale benchmark dataset and the K- and Radius-based Neighbors Intersection scores (KNI and RbNI) as metrics of model performance assessment on this benchmark that consider accuracy and alignment quality. Leveraging this benchmark, we find that the scVI architecture outperforms other methods in cross-dataset alignment. We enhance scVI’s performance by optimizing its architecture and show how adversarial learning can be used to boost performance further (BA-scVI). Optimizing architectures, loss functions, and parameters against metrics is common-place in machine-learning model development, for example (Kraus, Ba, and Frey 2016). With the release of the data and tests we develop here, we hope to increase adoption of these benchmark strategies in scRNA model development and spur development of yet better approaches.

It has been hypothesized that large cross-species atlases could be used to improve alignment quality (Lotfollahi et al. 2024). Supporting this, we find that aligning the large mouse scREF-mu atlas to the human scREF atlas improves accuracy on the mouse benchmark without reducing the quality of the human alignment or cell-type labeling. In an unbiased cell-type analysis of the aligned atlases, we identified complement-expressing macrophage and fibroblastic cell types that are prevalent across tissue types in both species. Peripheral complement expression has been described for macrophages (Müller, Hanauske-Abel, and Loos 1978). However, the same cannot be said for fibroblasts, and overall, complement expression has received limited attention (Bordron et al. 2020). We speculate that liver-expressed complement factors combine with those produced by macrophages in peripheral tissue and those of fibroblasts to modulate complement activity in humans and mice, pointing towards a broader underappreciated role in immunity (Figure 4k). In highlighting this result, we also hope to demonstrate the value of cross-species alignment and analysis for understanding cell-type evolution and conservation. We anticipate that such alignments at scale could be used to identify evolutionary relationships between cell types, thus tracing the emergence of the lineages we see today.

## Methods

### Collection and standardization of human scRNA data

Raw scRNA UMI count matrices were obtained from public repositories (Table S1). Quality control followed the original author filters. Cells labeled by the authors as; (i) Unknown; (ii) Undetermined; or (iii) Mixed were excluded from benchmark analysis. Gene identifiers were standardized across studies based on (i) Human Protein Atlas (HPA) versions 13 to 20; and (ii) ENSEMBL GRCh38 versions 78 to 103. Priority was given to HPA identifiers. Genes common to 30 datasets or more were used for training (Tables S1). In cases where the authors provided only general T-cell annotations, we used Azimuth’s Human PBMC signatures (Hao et al. 2021) to assign those cells into CD8+, CD4+ or gamma-delta T-cells.

### Collection and standardization of mouse scRNA data

Similar to human datasets, mouse data was obtained from public repositories (Table S1) and quality control followed the original author filters and cells labeled as unknown, undetermined or mixed were excluded from analysis. To align mouse and human datasets, gene names were uppercased and mouse-vs-human orthologs were mapped using ENSEMBL v110. Azimuth’s Human PBMC signatures were used to sub-classify general T-cells into CD8+, CD4+ or gamma-delta T-cells.

### scRNA data normalization

For scVI, PCAscmap, Harmony, and BA-scVI, the count matrices were normalized on a per-cell basis using Scanpy v1.7.2 (Wolf, Angerer, and Theis 2018), by dividing each cell by its total count over all genes. The normalized count was then multiplied by a scale factor of 10,000, after which a log(X+1) transformation was applied. For RPCA, Seurat’s SCTransform normalization was used with default parameters (Hao et al. 2021).

### Calculation of K-Neighbors Intersection (KNI) score

To calculate the KNI we consider the set *C* = {*c*_1_, *c*_2_ … *c*_*n*_ } of cells in a low-dimensional cell-type feature space where each cell *c*_*i*_ is defined by its coordinates *x*_*i*_, batch identifier *b*_*i*_, and cell-type identifier *t*_*i*_ . The distance function *D* between two cells is the Euclidean distance between their embedded coordinates. For the KNI, we thus identify the *k*-nearest neighbors for each cell *c*_*k*_ as per a K-nearest neighbors search:

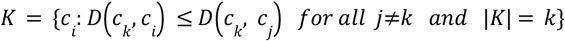

For each cell *c*_*k*_, we then identify a subset *B* of *K* in which cells have different batch identifiers, defined as:

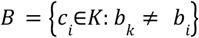

We then use a function L to assign ‘predicted’ cell-type labels to each datapoint. Each cell *c*_*k*_ is labeled as either (1) an outlier if the number of elements in *B* is below a threshold number τ (τ < *k*), i.e., too many nearest neighbors belong to the same batch, or (2) the most common label from cells in *B*:

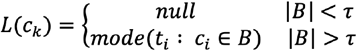

The KNI score is then calculated as the total number of predicted labels that match author given labels:

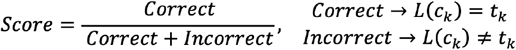

An analysis of KNI parameters is given in the supplemental. For comparisons performed in this paper, k=50 neighbors was used, with a cut-off τ of 40. For the model comparisons performed here quantile normalization was used (25%, 75%) to scale the embedding spaces prior to determination of the KNI. For optimization studies, the scikit-learn KNN algorithm was used (Pedregosa et al. 2012), while for scREF we use the FAISS GPU implementation of KNN search (Johnson, Douze, and Jégou 2021).

### Calculation of Radius-based Neighbors Intersection (RbNI) score

Calculation of the RbNN proceeds as per the KNI, except that: (1) The set of neighboring cells is defined by a radius *r*, as per Radius-based Nearest Neighbors:

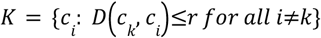

; (2) a threshold percent of ‘self’ data points τ^*^ is used; and (3) cells with no neighbors within the radius *r* are also given an outlier label:

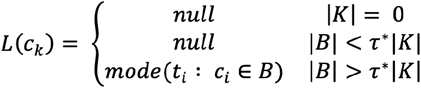

An analysis of RbNI parameters is given in the supplemental. For comparisons performed in this paper, a radius *r*=1.0 was used alongside a cut-off % of τ*=80%, alongside the same quantile normalization as performed for the KNI. For the smaller scMARK benchmark we use for optimization (Suplemental), the scikit-learn Radius-based nearest neighbors algorithm was used (Pedregosa et al. 2012) while for scREF we use the FAISS GPU implementation of Radius based search (Johnson, Douze, and Jégou 2021).

### Cell-type space alignment parameters

Methods were implemented as follows:

- **PCA**: Highly variable genes were selected as relevant features on the basis of higher dispersion than genes with similar mean expression (Satija et al. 2015), as implemented in *scanpy* v1.7.2 (Wolf, Angerer, and Theis 2018). PCA was run on the scaled, normalized expression of those highly variable genes,
- **RPCA:** The RPCA method was implemented in R using Seurat (v4.0.3) (Hao et al. 2021) with the top-10 larger samples used as references for anchor detection and the following parameters (dims=10, npcs=10, k.filter=150, k.weight=100). The output of the functions RunPCA (npcs=10) and RunUMAP (n.components=10), with assay=“SCT” were used as inputs for KNI or RbNI calculations.
- **Harmony:** The PCs identified from highly variable genes with PCA as described above were passed to a python implementation of Harmony, *harmony-pytorch* v.0.1.7, using default parameters (https://doi.org/10.1038/s41592-019-0619-0).
- **scVI 2L Sample:** We reimplemented the scVI variational auto-encoder described by (Lopez et al. 2018). We used sample level batch-correction with the following hyper-parameters: (1) 2-layer encoder and decoders; (2) 512 hidden nodes for each linear layer; (3) Dropout regularization with 0.1 probability of an element to be zeroed; (4) Batch normalization in between two hidden layers; and (5) The ReLU activation function. The latent space dimension was set to 10 and modeled using a Normal distribution. A Zero-Inflated Negative Binomial (ZINB) distribution was used to model gene counts as per (Lopez et al. 2018). The Adam optimizer was used for training the VAE with learning rate = 1E-4; weight decay = 1E-5; and eps = 0.01 (Kingma and Ba 2014). Early stopping was used with patience = 15 epochs. The model was trained with a batch-size 64 and for a maximum of 100 epochs. All the implementation was done in Python using Pytorch (1.7.0) library. One-hot Batch ID vectors corresponded to the unique Sample ID, 386 batches/samples were defined over the 10 studies.
- **scVI 4L Sample:** An optimized scVI model was identified (Table S3) based on parameter and architecture optimisations. This model was implemented as above, but leveraged 4 layers in the encoder and decoder (4L), and patience = 5 (lp) for the early stopping criteria.
- **scVI* (ScVI 4L-NoL-NoB Both) :** An optimized scVI model, *that did not require batch ID in the encoder*, was identified (Table S3) based on parameter and architecture optimisations outlined in the Supplementary for us on held-out data. This model was implemented as above, however; (1) Explicit handling of library size was removed (Lopez et al. 2018), (NoL); (2) *The batch ID vector was not injected into the encoder layer*, (NoB); (3) A two-hot batch ID vector was used that encoded ‘Both’ Sample ID (386 long), and study ID (11 long); and (4) learning rate = 5E-5 was used.
- **scGPT:** For both zero-shot and fine-tuned embeddings, where applicable we used the authors’ tutorials (accessed March 25th), 2024 for embedding. as well as preprocessing pipelines as described in the paper. https://github.com/bowang-lab/scGPT/blob/main/tutorials/zero-shot/Tutorial_ZeroShot_Integration.ipynb https://github.com/bowang-lab/scGPT/blob/main/tutorials/Tutorial_Integration.ipynb Models were retrieved from: https://drive.google.com/drive/folders/1oWh_-ZRdhtoGQ2Fw24HP41FgLoomVo-y (Cui et al. 2024) The 512-dimensional embeddings obtained from the fine-tuned scGPT models were reduced to 10 dimensions using a two layer autoencoder trained using cosine similarity loss. This step enabled direct comparison with the 10-dimensional BAscVI embeddings.
- **geneFormer:** We used the authors’ provided zero-shot pipeline accessed March 27th, 2024) for preprocessing, tokenization, and embedding. https://huggingface.co/ctheodoris/Geneformer/blob/main/examples/tokenizing_scRNAseq_data.ipynb https://huggingface.co/ctheodoris/Geneformer/blob/main/examples/extract_and_plot_cell_embeddings.ipynb We extracted embeddings using the provided weights (Theodoris et al. 2023), retrieved from: https://huggingface.co/ctheodoris/Geneformer/tree/main/geneformer-12L-30M For both the geneFormer and scGPT embedding procedures dataloaders leveraging the TileDB database were used (TileDB, 2024), while for VAE models (scVI, BA-scVI) data-loaders loaded data in directly from H5AD files. All data-loaders and model training procedures leveraged the PyTorch lightning library.

### BA-scVI architecture

Batch-Adversarial scVI (BA-scVI) leverages the same core architecture as scVI, but makes use of an adversarial framework for removing batch effects. The key difference is where scVI injects one-hot batch ID vectors into the encoder and decoder layers, BA-ScVI takes an adversarial learning approach to learning and removing batch-effects.

- Here discriminators seek to predict the batch-ID *b*_*i*_ using the encoder outputs and decoder inputs. Namely, the discriminator *D* seeks to minimize loss *L* with respect to batch-ID on the encoder outputs *W*_*E*_ and decoder outputs *W*_*D*_ . The encoder and decoder weights are frozen in this step. We use cross entropy loss such that,

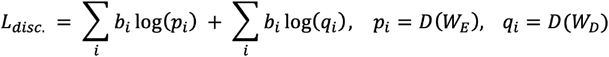
- The inference network then seeks to: (1) Maximize the probability of the posterior, which in this case we use a Zero-Inflated Negative Binomial (ZINB) distribution as per ((Lopez et al. 2018); (2) Minimize KL-divergence of the embedding distribution *z* and library encoder *l* (Kingma and Welling 2013); and (3) Maximize discriminator loss, i.e.,

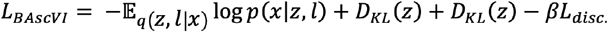

The discriminator and inference networks are then trained in sequential steps with the first step used to update weights on the discriminator networks and the second step weights on the inference network. An optimal regimen for training was identified (Table S3) that leveraged an Adam optimizer (Kingma and Ba 2014), with learning rate = 5E-5 for the inference network, 1E-2 for the discriminator network; weight decay = 1E-5; and eps = 0.01, with a batch-size 64 and for a maximum of 100 epochs; β = 10 was used for the model described in the main text. Other values of beta were tested, but performed similarly or worse than the top performing model that emerged from initial optimization studies (Table S3). In this optimal training regime a two-hot batch ID vector was also used that encoded ‘Both’ Sample ID (386 long), and study ID (11 long) was also used.

### Model training on scREF, scREF-mu and the joint scREF/scREF-mu atlas

Models were trained on scREF, the scREF/scREF-mu atlas using a regime optimized on a smaller benchmark scMARK that we discuss in the supplemental, with the exception of our handling of a standardized gene set for training. For scMARK genes common to all datasets were used. For scREF and the joint atlas we took a list of genes common across 30 datasets or more. To handle missing genes for a specific dataset, we then applied a mask to the reconstruction loss function at train time, such that only genes present in the dataset affected the overall loss. This mask was not applied to either the encoder or decoder, and thus will not affect prediction results. For the joint atlas, we used ENSEMBL v110 (Martin et al. 2023) . On scREF-mu, mouse genes identifiers common to all datasets were used (Table S1).

### Comparison of label propagation and K-means methods for cell-type ID

Label propagation was implemented as described in (Raghavan, Albert, and Kumara 2007). KNN graphs for Label propagation were generated using the FAISS library (Johnson, Douze, and Jégou 2021), setting the number of neighbors to k=50. Label propagation was run until complete convergence, with the subsequent number of clusters identified by label propagation used to define the number of clusters *k* for k-mean clustering. The sci-kit Learn K-means clustering function was used for clustering (Pedregosa et al. 2012), with 150 iterations set as the maximum number of iterations. The Adjusted Rand Index was calculated using the function provided in SciPy version 1.13.0.

### Differential Gene Expression

To identify enriched genes in the unbiased cell-type clusters we calculated differentially expressed genes between the cell-type cluster and the distribution of genes across all cell-types including that cluster, given that the size of the dataset was many times larger than the largest cell-type grouping. This ensured that results would not be distorted by including the group in question in the reference cluster, and the number of calculations run was dramatically reduced. To enable us to store a full Scanpy annotated data object in memory (Wolf, Angerer, and Theis 2018) for DGE calculations (64GB RAM), we took a sample of 2,000 cells within a cell-type cluster, or all cells in the cluster (whichever value is smaller). Samples were taken from a TileDB array (TileDB, 2024)) of the full scREF object that we make available. Raw read-counts were standardized as described in (Stuart et al. 2019), and we filtered genes with very low expression (total standardized read-count <20 across all cell-types). To then calculate differentially expressed genes, we calculated Welch’s T-test score between the cell-type grouping distribution and the distribution across the full cell sample. We multiplied the reference standard deviation for each gene by a factor of 10 to enhance the ranking of genes that show large effect size differences between the cluster and reference groups. The top 25 genes for each cell-type cluster were assessed visually and passed into ShinyGO 0.8 for pathway enrichment analysis (Ge, Jung, and Yao 2020). We provide the top 250 genes by DGE in Table S5 for each cluster, alongside the cluster ID, and cell-type composition of the cluster and sample based on standardized and author cell-type labels and tissue types represented in the cluster. A round of hierarchical clustering on cluster gene expression means was also performed using scikit-learn on the cluster medians to provide an easy to explore ordering of the clusters in the Tables.

We also provide the top 250 genes by DGE for a more complete set of 591 human filtered for at least 30 cells present in at least 2 studies, 368 mouse clusters filtered for at least 20 cells present in at least 2 studies, for the benefit of the reader in Supplemental Table S5. Here samples of 250 cells were taken for each human cell-type grouping and 300 for mise. We note visual inspection suggested no clear differences in top differentially expressed genes occur between 250 cell and 2,000 cell samples indicating these are sufficient sample sizes for ID of top differentially expressed genes. Again cluster composition metrics are provided and ordering is as per hierarchical clustering of cluster gene expression medians.

### Heatmaps of Complement Pathway expression

Raw read-counts were standardized as described in (Stuart et al. 2019). Heatmaps of complement pathway expression were calculated by then taking the mean across all cells of a specific author standardized cell-type and tissue-type.

#### Filtering of artifactual clusters of conserved cell-types

Alongside filtering of clusters identified by LP for those present in at least 10 tissue-types, we also removed those groupings that were highly enriched for mitochondrial or ribosomal genes as determined by differential expression. Specifically, those

#### UMAP projections

All UMAP projections were produced using the UMAP library version 0.5.6 with default setting unless otherwise specified (McInnes, Healy, and Melville 2018). For the online tool we provide, UMAP is calculated using the parametric version of UMAP described in Sainburg et al. (Sainburg, McInnes, and Gentner 2021).

## Contributions

S. Singh and O. Focsa implemented and ran initial testing of scVI and built the pipeline for model testing and optimization. D. Mendonca contributed to the ML pipeline testing, built the web-interface for visualization of alignment results, and helped with the manuscript. M. Buechler helped in writing the manuscript and guiding the direction of the paper. A. Nixon, R. Schuster, B. Hinz generated the in vitro scRNA data used for alignment. E. Williams ran the embedding pipeline for scGPT and Geneformer. B. Innes benchmarked the Harmony and PCA models. J. J. Diaz Mejia collected, curated, and standardized the scMARK, scREF, and scREF-mu datasets, and contributed to writing the manuscript. S. Cooper conceived the work, ran the model and metric optimization studies for the scVI and BA-scVI models, analyzed the aligned atlases, and wrote the manuscript. We would like to acknowledge Sean Grullon for help conceiving the project and Sarah Hackett for help curating scREF metadata. We are grateful for the UHN STARR facility for helping generate the *in vitro* scRNA sequencing data. We would like to thank Christoph Licht, Mike Briskin, Christopher Harvey, Jimmy Ba and Girish Aakalu for critical comments, revisions, and help on the manuscript. Finally, we would like to acknowledge the rest of the team at Phenomic for feedback on the project generally.

## Data and Code Availability

The smaller scMARK benchmark we provide is publicly available at: https://zenodo.org/record/7795653, The larger scREF and scREF-mu dataset available as TileDB objects at https://cloud.tiledb.com/ under the “public/phenomic” repository - we can provide links to the files in ‘.mtx’ format on request.

Functions used to access our scRNA embedding capability programmatically, alongside code and configuration files used to train all models and neural networks, links to weight for trained models, and code used to generate key figures is all available at:

https://github.com/samocooper/bascvi

UMAP projections, dotplots of the alignments, DGE, and drag-and drop scRNA alignment capabilities can all be interactively explored at:

https://scref.phenomic.ai/

## Supplementary Data

Supplemental: Development Of the KNI and RbNI metrics for cell-type alignment evaluation

Table S1: List of studies used in the scMARK and scREF datasets, cell-type mappings for standardization, genes used for training, barcodes used in final score calculation

Table S2: Agreement of RbNI and KNI scores with other scores across experiments

Table S3: Optimization of models against smaller scMARK benchmark dataset

Table S4: Evaluation of model performance on the scREF dataset using KNI / RbNI scores

Table S5: Evaluation of model performance on the scREF-mu dataset using KNI / RbNI scores

Table S6: Analysis of label propagation groupings by differential expression on scREF and scERF-mu

