## Supplemental Material for "Adversarial learning enables unbiased organism-wide cross-species alignment of single-cell RNA data at scale"

### Supplemental Information to "Adversarial learning enables unbiased cross-species single-cell RNA atlas alignment and reveals conserved cell-types between mice and humans"

##### *Authors*

Juan Javier Díaz-Mejía[1], Elias Williams[1], Brendan Innes[1], Octavian Focsa[1], Dylan Mendonca[1], Swechha Singh[1], Allison Nixon[1], Ronen Schuster[1,2], Matt Buechler[3], Boris Hinz[2,4], and Sam Cooper[1]

##### *Affiliations*

1. Phenomic AI Inc., MaRS Centre, West Tower, 661 University Ave Suite 1300, Toronto, ON M5G 0B7
2. Faculty of Dentistry, University of Toronto, Toronto, M5S 3E2 Ontario, Canada
3. Laboratory of Tissue Repair and Regeneration, Keenan Research Centre for Biomedical Science of the St. Michael's Hospital, Toronto, Ontario M5B 1T8, Canada

##### *Metric Evaluation and development of Radius-based Neighbors Intersection Score*

In this study we make the initial assumption that on average Author labels are correct. Thus, in good cell-type spaces, cells with the same label will align, while in bad cell-type spaces, batch effects will dominate. In this supplemental we consider two simulated examples, one simple, the other more realistic, to arrive at metrics we term the K-neighbors and Radius-based Neighbors Intersection (KNI and RbNI respectively) scores as effective and scalable means of evaluating the quality of an aligned cell-type space given a set of ground-truth author labels. Later we compare how the set of scores evaluated rank the models considered in the main-text on a simple test case; this further supports the KNI and RbNI scores as effective metrics for evaluating cell-type space alignment.

##### *Analysis of metric behaviors on a simple simulated test case*

In the first example we consider the theoretical case of two cell-types $C_1$, $C_2$ and batches $W_1$, $W_2$. Each cell in this analysis has both a cell-type and a batch it corresponds to. The probability distribution of a given cell's type $C$ (either $C_1$ or $C_2$) and the probability distribution of a given cell's batch $W$ (either $W_1$ or $W_2$) can each be approximated using a Gaussian mixture model with two components. The cell type probability distribution is therefore the sum of two normal distributions with the same variance, where mean values are offset by φ. Similarly, the batch probability distribution is the sum of two normal distributions with the same variance, where the mean values are offset by ω instead.

$$X \sim \begin{bmatrix} C \\ W \end{bmatrix} = \begin{bmatrix} N(\mu, \sigma^2) + N(\mu + \psi, \sigma^2) \\ N(\mu, \sigma^2) + N(\mu + \omega, \sigma^2) \end{bmatrix}$$

In (Figure S1a) each point on the graph is a cell that has a cell-type and belongs to a batch. A random sampling of a cell, $X$, therefore is represented by a 2D vector where the first dimension corresponds to the cell-type probability distribution and the second dimension is the batch probability distribution.In a good cell-type space, cells are well separated ($\varphi \gg 0$), meaning a vertical line can discriminate cell types, while batch effects are small ($\omega \approx 0$), meaning no horizontal line can discriminate between batches. We therefore are seeking a metric which is large when $\varphi \gg 0,\ \omega \approx 0$. In a bad cell-type space cells are poorly separated ($\varphi \approx 0$), while batch effects are large ($\omega \gg 0$). Thus we are also seeking a metric which is small when $\varphi \approx 0,\ \omega \gg 0$.

We empirically compared 6 metrics for evaluating this theoretical cell-type space on three test cases, generated by sampling 2000 cells, split 50:50 between cell-types and batch effects; (1) An 'ideal' cell-type space where $\varphi = 4.0,\ \omega = 0$; (2) A case with poor cell-type separation but no batch effects $\varphi = 2.0,\ \omega = 0$; and (3) A case with good cell-type separation but large batch effects $\varphi = 4.0,\ \omega = 2.0$ (Figure S1a).

The metrics tested were; (1) Accuracy of a Bisecting line in the cell-type axis at $\varphi/2$ (Bis.) (2) Accuracy of a Support Vector Machine, trained to predict $C_1$ vs. $C_2$ on $W_1$ and tested on $W_2$ (SVM); (3) Accuracy of K-Nearest Neighbors model at predicting $C_1$ vs. $C_2$ fit on $W_1$ and tested on $W_2$ (KNN; k=25); (4) The Silhouette coefficient calculated pairwise between the cell-type clusters $C_1$ and $C_2$ (Sil.); (5) Mutual Information between 2 cluster labels assigned by a K-means clustering algorithm, and the cell-type cluster labels $C_1$and $C_2$ (KmMI); and (6) Accuracy of Radius-based Nearest Neighbors to predict $C_1$vs. $C_2$ fit on $W_1$ and tested on $W_2$; here points in $W_2$ that are beyond a certain radius $r$ to any point in $W_1$ are given an outlier label. The results are given in Figure S1a (below). Model parameters are scikit-learn defaults unless specified (Pedregosa et al. 2011).

In this analysis we see that when clusters are moved apart in the orthogonal direction, i.e. $\omega > 0$, only the Silhouette score and Radius-based Nearest Neighbors (RbNN) score drop in value (Figure S1a, No Batch, Sil.; $0.558 \rightarrow 0.498$; No Batch, RbNN, $0.816 \rightarrow 0.447$). The incorporation of the bisecting line highlights why this is the case since the SVM, KNN, and K-means search algorithms all approximate this partition. The Silhouette score declines since the intra-cluster distance reduces in the orthogonal direction (Rousseeuw 1987). In the RbNN approach, an outlier label is specified for points that have no neighbors within a certain radius in the training dataset. The RbNN score declines since a radius $r$ is chosen that is below the separation distance $r < \omega$. Thus, a number of points are labeled as outliers when $\omega = 2$. This simple example thus highlights the need for more explicitly penalizing batch effects in evaluation metrics, that would otherwise go undetected.

We thus constructed a set of 6 modified evaluation metrics that penalize batch effects more directly, and evaluated the performance of these methods on the three tests cases;

1. ***Bisecting Line (Bis.):*** We modified the bisecting line by including a second bisecting line in the batch axis at $\omega/2$, the accuracy of this line at predicting batch is deducted from the cell-type prediction accuracy of the 1st bisecting line in the cell-type axis at $\varphi/2$. Here we find that the bisection score is now substantially less in the case of $\varphi = 4, \ \omega = 2$ than it is in $\varphi = 4, \ \omega = 0$.

2. ***Silhouette coefficient Line (Sil.):*** We adjusted the the Silhouette coefficient to further penalize batch separation by deducting the coefficient calculated between batch-clusters from cell-type clusters i.e., $SC(C) - SC(W)$. The resulting score decreased much further when batches were separated (Figure S1a, Batch, SVM, $0.557 \rightarrow 0.341$). A similar decrease was seen when cell-type clusters were closer.

3. ***Support Vector Machine (SVM):*** We adjusted the SVM by deducting the accuracy of predicting batch from the accuracy of predicting cell-type. In predicting batch accuracy we trained on one cell-type and tested on the other, in predicting cell-type accuracy we trained on one batch and predicted on the other; thus 'cross validating in two axes'. This adjusted SVM score severely decreased on the addition of batch effects (Figure S1a, Batch, SVM, $0.473 \rightarrow 0.129$).

4. ***K-means Mutual Information (KmMI):*** In the KmMI approach, we increased the number of clusters $K$ to 4 to account for all membership possibilities $x \in C_i, W_j, (i, j) : \{0, 1\}$. We then deducted MI calculated between the 4 cluster labels vs. ground-truth batch labels, from the MI calculated between the 4 cluster labels vs. ground-truth cell-type labels i.e., $MI(K; C) - MI(K; W)$. This modified KmMI fell significantly when batch effects were introduced (Figure S1a, Batch, KmMI, 0.589 $\rightarrow$0.315).

5. ***K-Neighbors Intersection (KNI):*** When looking to extend KNN to penalize batch effects, we identified an approximation that significantly simplified the complexity of calculating the KNN score as defined earlier - and incorporates the KBET score as defined in (Büttner et al. 2019).

To calculate the KNI we consider the set $C = \{c_1, c_2 ... c_n\}$ of cells in a low-dimensional cell-type feature space where each cell $c_i$ is defined by its coordinates $x_i$, batch identifier $b_i$, and cell-type identifier $t_i$. The distance function $D$ between two cells is the Euclidean distance between their embedded coordinates. For the KNI, we thus identify the $k$-nearest neighbors for each cell $c_k$ as per a K-nearest neighbors search:

$$K = \{c_i : D(c_k, c_i) \leq D(c_k, c_j) \ for\ all\ j \neq k \ and\ |K| = k\}$$

For each cell $c_k$, we then identify a subset $B$ of $K$ in which cells have different batch identifiers, defined as:

$$B = \{c_i \in K : b_k \neq b_i\}$$

Each cell $c_k$ is then labeled as either (1) an outlier if the number of elements in $B$ is below a threshold number $\tau$ $(\tau < k)$ , i.e., too many nearest neighbors belong to the same batch, or (2) the most common label from cells in $B$.

$$L(c_k) = \begin{cases} null & |B| < \tau \\ mode(t_i : c_i \in B) & |B| > \tau \end{cases}$$

The KNI score is then calculated as the total number of predicted labels that match author given labels:

$$Score = \frac{Correct}{Correct + Incorrect}, \quad \begin{matrix} Correct \rightarrow L(c_k) = t_k \\ Incorrect \rightarrow L(c_k) \neq t_k \end{matrix}$$

We term this metric the K-nearest Neighbor Intersection (KNI) score since it only evaluates membership at the intersection of clusters (cell-types) in the dataset (Figure S1b). We note generalizations of this metric could require intersection with multiple clusters.

Increasing the number of $k$-nearest neighbors tested increases the size of the search around the intersection, while increasing the cut-off $\tau$ increases the threshold for the number of neighbors that need to be from the same batch for the data-point to be assigned a null-value (Figure S1b). In the case of perfect alignment when $\varphi = 0$, $\omega = 0$, we find that when $k = 25$ and $\tau = 20$, 100% of data points are given a label demonstrating for suitable $\tau$ the KNI approaches 1.0 in the perfect case. At $\tau = 15$ ~80% of data points are labeled, indicating that higher $\tau$ values are needed to ensure areas of significant overlap are given labels based on their K-nearest neighbors. With the KNI we now see a substantial score reduction when datasets are moved apart in the batch direction (Figure S1a, Batch, KNI; $0.981 \rightarrow 0.365$)

6. ***Radius-based Neighbors Intersection (RbNI):*** Similarly, we determined that we could modify RbNN to include an outlier label if too many values within a certain radius are 'self'; we term this Radius-based Neighbors Intersection (RbNI), and calculation of the RbNN proceeds as per the KNI, except that: (1) The set of neighboring cells is defined by a radius, as per Radius-based Nearest Neighbors such that,

$$K = \{c_i : D(c_k, c_i) \leq r \; for \; all \; i \neq k\}$$

; (2) a threshold percent of 'self' data points $\tau^*$ is used; and (3) cells with no neighbors within the radius $r$ are also given an outlier label such that,

$$L(c_k) = \begin{cases} null & |K| = 0 \\ null & |B| < \tau^*|K| \\ mode(t_i : c_i \in B) & |B| > \tau^*|K| \end{cases}$$

The RbNI also shows a substantial score reduction when datasets are moved apart in the batch direction (Figure S1a, Batch, RbNI, $0.895 \rightarrow 0.381$). These metrics also maintain high scores in ideal scenarios (close to 1 in this example), since the null accuracy of predicting batch is not deduced.

Overall, these adjusted metrics thus make intuitive sense for evaluating cell-type spaces where the goal is to promote cell-type alignment while penalizing batch effects. However, we noted that calculating the SVM metric is complex due to the need for cross validating on both held-out 'cell-types' and batches. Realizing the complexity of this cross validation is magnified when 'cell-types' are poorly defined we thus dropped the SVM metrics. For example, on real data, should cross validation occur over labeled cell-types, or random 10% splits across all cells?

The RbNN, KmMI, KNI, and RbNI metrics demonstrate potential for assessing cell-type space alignments, but all require selection of specific parameters, which could lead to bias in using these metrics for evaluating cell-type spaces. We thus sought to understand on this theoretical case how these models score the 'perfect' alignment case of $\varphi = 4, \; \omega = 0$ vs. the batch effect case $\varphi = 4, \; \omega = 2$ while varying metrics parameters.

**Figure S1 (Below): Analysis of metric behavior on theoretical example:** a) Scatterplots of the three test cases used to compare candidate metrics for evaluating aligned metrics, the key parameters are separation of the two cell-types by φ and separation of the two batch effects by ω. Metric scores for these three test cases are provided below, and are separated by those that are not adjusted to account for batch effects, and those that are; b) Under the KNI score, cells that are surrounded by more than τ cells from the same batch are classified as null, this example demonstrates the effect of this, by considering two batches separated by a batch effect ω. Cells that are labeled as null are blue, vs. those that would be tested for label accuracy red, two regimes are considered τ = 15 and τ = 20, showing the effect on the KNI score in the case of large batch effects ω = 4, and no batch effect ω = 0; c) The RbNN score on the ideal case ($\varphi = 4, \; \omega = 0$) vs. the case of batch effects ($\varphi = 4, \; \omega = 2$) as plotted in (a) while varying the parameter radius *r*; d) The same plot as (c) for the KmMI score where the cluster number parameters is varied; e) The separation between the scores for the two test cases, varying *r* for the RbNN (Orange) and cluster number for the KmMI (blue); f) same as prior examples but considering the KNI score, and varying the cutoff parameter τ and number of nearest neighbors k, the score difference between the two test cases for the two parameters is also plotted (left); g) same as (f) but for the RbNI, where the cutoff percent $\tau^*$ and radius parameters $r$ are varied.

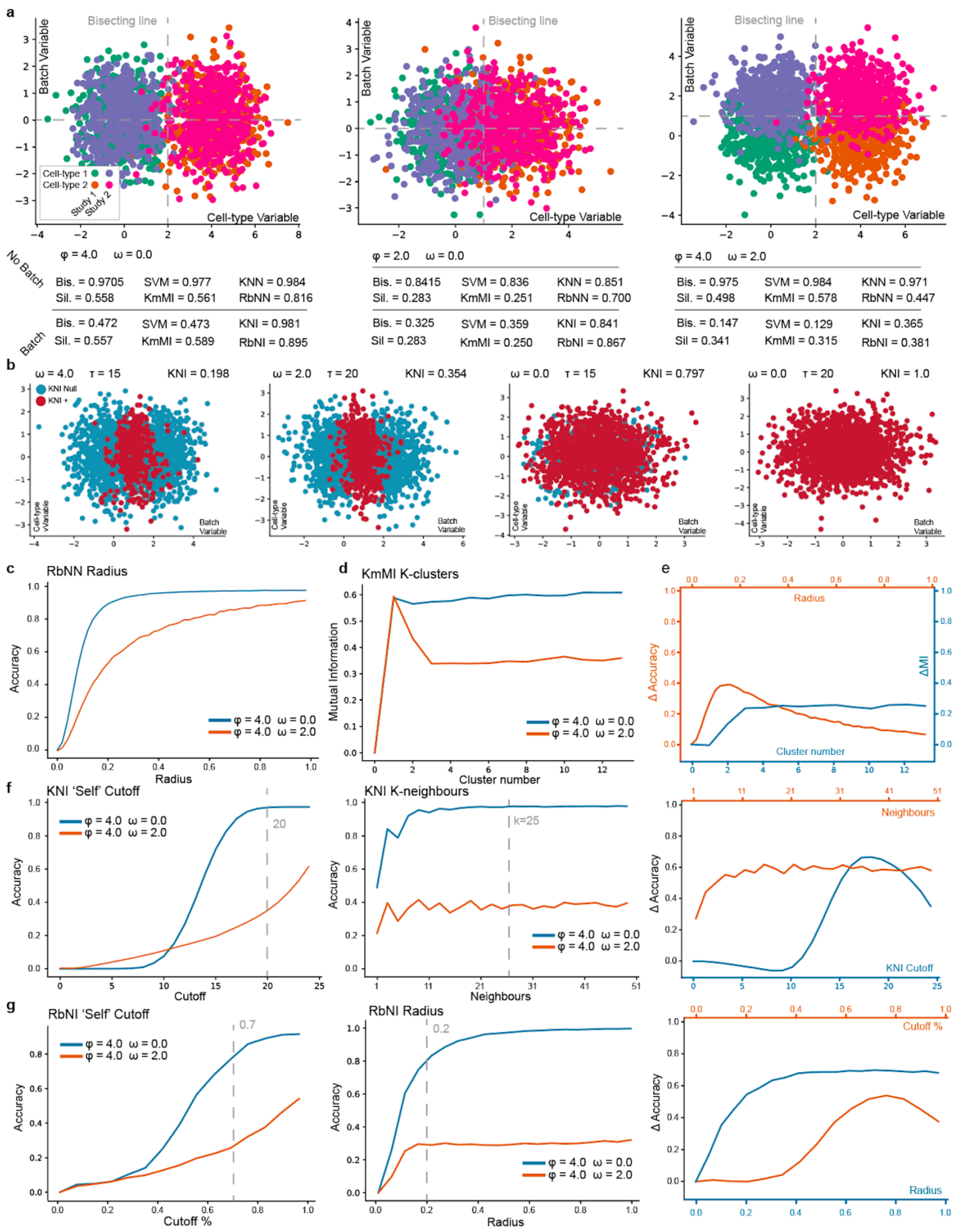
a
Bisecting line
Batch Variable
Cell-type Variable
Cell-type 1
Cell-type 2
Study 1
Study 2
No Batch
Batch
φ = 4.0 ω = 0.0
Bis. = 0.9705 SVM = 0.977 KNN = 0.984
Sil. = 0.558 KmMI = 0.561 RbNN = 0.816
Bis. = 0.472 SVM = 0.473 KNI = 0.981
Sil. = 0.557 KmMI = 0.589 RbNI = 0.895
φ = 2.0 ω = 0.0
Bis. = 0.8415 SVM = 0.836 KNN = 0.851
Sil. = 0.283 KmMI = 0.251 RbNN = 0.700
Bis. = 0.325 SVM = 0.359 KNI = 0.841
Sil = 0.283 KmMI = 0.250 RbNI = 0.867
φ = 4.0 ω = 2.0
Bis. = 0.975 SVM = 0.984 KNN = 0.971
Sil. = 0.498 KmMI = 0.578 RbNN = 0.447
Bis. = 0.147 SVM = 0.129 KNI = 0.365
Sil = 0.341 KmMI = 0.315 RbNI = 0.381
b
ω = 4.0 τ = 15 KNI = 0.198
ω = 2.0 τ = 20 KNI = 0.354
ω = 0.0 τ = 15 KNI = 0.797
ω = 0.0 τ = 20 KNI = 1.0
KNI Null
KNI +
Cell-type Variable
Batch Variable
c
RbNN Radius
Accuracy
Radius
φ = 4.0 ω = 0.0
φ = 4.0 ω = 2.0
d
KmMI K-clusters
Mutual Information
Cluster number
e
Radius
Δ Accuracy
ΔMI
Cluster number
f
KNI 'Self' Cutoff
Cutoff
20
KNI K-neighbours
Neighbours
k=25
KNI Cutoff
g
RbNI 'Self' Cutoff
Cutoff %
0.7
RbNI Radius
0.2
Cutoff %

1. ***RbNN:*** For the RbNN we varied the search radius $r$. In this example, the most significant separation between the ideal and batch effect cases exists at a radius of ~0.17 (Figure S1c, e). For high $r$ the differences between the two cases tailed off considerably, indicating batch effects can only be seen over a small range of $r$.

2. ***KmMI:*** The KmMI metric was most robust to variation in the key parameter selection of $k - clusters$. Indeed, beyond a cluster number of 3, almost no change in the MI value is seen (Figure S1d, e). Thus, both the RbNN and KmMI approaches could work although the RbNN metric is likely only sensitive over a narrow range of $r$; this could be difficult to identify.

3. ***KNI:*** The KNI has two adjustable parameters, the number of neighbors $k$ and the cut-off for the number of neighbors labeled as 'self' $\tau$. First, we fixed the number of neighbors to analyze with the KNI metric to $k = 25$. Then varying the KNI metric $\tau$ showed a peak separation at $\tau = 17$. For values below $\tau = 17$ the accuracy score on the ideal case $\varphi = 4$, $\omega = 0$ also dropped, due to the effect of too many 'self' data-points being close, as per Figure S1f. Importantly, for a wide range of parameter values $15 < \tau < 24$ the KNI score shows a large separation between the ideal and batch-effect scenario. We then adjusted the number of neighbors $k$ used in KNI, under a fixed cutoff of 80% (i.e. 20 when $k$=25). This had a minimal impact on separation for all $k > 5$. Indicating the KNI is sensitive to the choice of cutoff value below a certain level, but insensitive to varying the number of neighbors analyzed.

4. ***RbNI:*** The RbNI has two adjustable parameters, the number of neighbors within a radius $r$ and the cut-off for the number of neighbors labeled as 'self' $\tau^*$. Here, varying $\tau^*$ had a significant impact on the score but, similar to the KNI cutoff $\tau$, for a wide range of parameter values $60\% < \tau^* < 100\%$ the RbNI score shows a large separation between the ideal and batch-effect scenario Figure S1g. Meanwhile, varying the radius parameter beyond a small value, e.g., 0.2 had no effect. This shows that for the RbNI there also exists parameters where a high degree of separation between key test cases can be seen.

In summary, Batch Adjusted Silhouette score, RbNN, KmMI, KNI, and RbNI, all demonstrate potential for being able to evaluate cell-type space alignments based on this simple theoretical test case. Where the silhouette score has no parameters that can be varied, the RbNN, KmMI, KNI, and RbNI all do. We note that while the RbNN score is sensitive to choice of *r* the other metrics are robust over a wide range of parameters.

##### *Analysis of metric behavior on cell-type space from one dataset with spiked noise/batch-effects*

In this second example, we sought to understand how the Batch Adjusted Silhouette score, RbNN, KmMI, KNI, and RbNI perform on a more realistic simulated example of an aligned scRNA cell-type space $S$. To create this more realistic example we took a 30,000 cell sample of a single high-quality scRNA dataset (Bassez et al. 2021) and embedded it into a 5-dimensional space $S \in R^5$ using a basic implementation of a Variational Autoencoder with Mean Squared Error loss function (Kingma and Welling 2013), (VAE MSE; 2 layer encoder and decoder, 512 neurons per hidden layer, lr = 1E-4, patience = 15). We chose this model so as to reduce bias in our model comparison, where we anticipate published models specifically designed for scRNA analysis should outperform this approach. We took the resulting embedded cell-type space and simulated both poorer alignment, and worse batch-effects, to compare how the metrics scores these spaces vs. the unmodified VAE encoding.

***Alignment:*** To simulate progressively worse alignment, we introduced random gaussian noise to the cell-type space $S + N(\mu, \sigma^2)$ where $\mu = 0$ and we vary $\sigma^2 = \{0, 0.2, 0.4, 0.6, 0.8\}$. Data was then re-normalized to unit mean and variance after the addition of noise. UMAP projections of the cell-type spaces for cases $\sigma^2 = \{0, 0.4, 0.8\}$ are given in Figure S2a and highlight the progressive increase in cell-type cluster overlap associated with introduction of this noise. We thus sought to assess how the metrics scored these increasingly noisy cell-type spaces, while varying metric parameter space.

1. ***Batch Adjusted Silhouette Score:*** Analyzing the Silhouette score calculated on the ground-truth cell-type label clustering indicated that this value decreased approximately linearly, from 0.0267 through to -0.0591 . These values were much smaller than those calculated for the theoretical case of two batch effects and two cell-types, likely a result of higher dimensionality and many more clusters. The fact that this score decreases with increasing noise indicates it may still be effective in cell-type space evaluation.

2. ***RbNN:*** The RbNN metric is able to separate good from bad cell-type embedding spaces over a wide range of $r$ in this test, though we note at small values of $r$ it is only sensitive to small additions of noise, while at larger values, $r > 0.3$, sensitivity to the addition of noise drops off quickly (Figure S2b). Of note, the RbNN approach is non-linear with respect to the amount of noise added, with the initial noise, $\sigma^2 = \{0, 0.2\}$ reducing the score much more than subsequent noise, e.g., $\sigma^2 = \{0.6, 0.8\}$. This trait is valuable since valuable small improvements to top performing models will be better separated.

3. ***KmMI:*** the KmMI score showed a linear relationship to the addition of noise, and was only able to separate the smallest addition of noise $\sigma^2 = \{0.2\}$ at very large cluster numbers $\sim k > 100$. However, at these higher cluster numbers robust separation between all scenarios was observed (Figure S2c). Thus while the KmMI approach is likely less sensitive to small variations in cell-type alignment space, at high cluster numbers it may represent a valuable tool for evaluating cell-type alignment space, and benefits from only having a single tunable parameter.

4. ***KNI:*** The KNI score showed a linear decrease in score for all cut-off values greater than $\tau = 17$ indicating this score is effective over a reasonably wide set of parameters (Figure S2d). Of note, this is the same value of $\tau$ for which optimal separation of scores was seen in the theoretical case described in the first section, indicating this parameter may also be robust to dimensionality and dataset complexity. Varying the number of neighbors $k$ while setting $\tau$ as 75% of the value of $k$ also highlights that for values of $k > 10$, robust separation between all scenarios is observed.

5. ***RbNI:*** Finally, the RbNI score showed a similar nonlinear sensitivity to the RbNN score with respect to the addition of noise, a valuable property as noted above. Similar to the KNI the RbNI score was insensitive to

the selection of the cutoff percent τ*, for τ>50% when the radius was fixed at a value of 0.3, indicating stability here (Figure S2e). This result also matched that seen in the theoretical test case above, suggesting that this parameter is likely stable of cell-type space dimensionality and complexity. Finally a similar relationship between selection of *r* and sensitivity to addition of noise was seen between the RbNI and RbNN methods, when a cutoff of 75% was used.

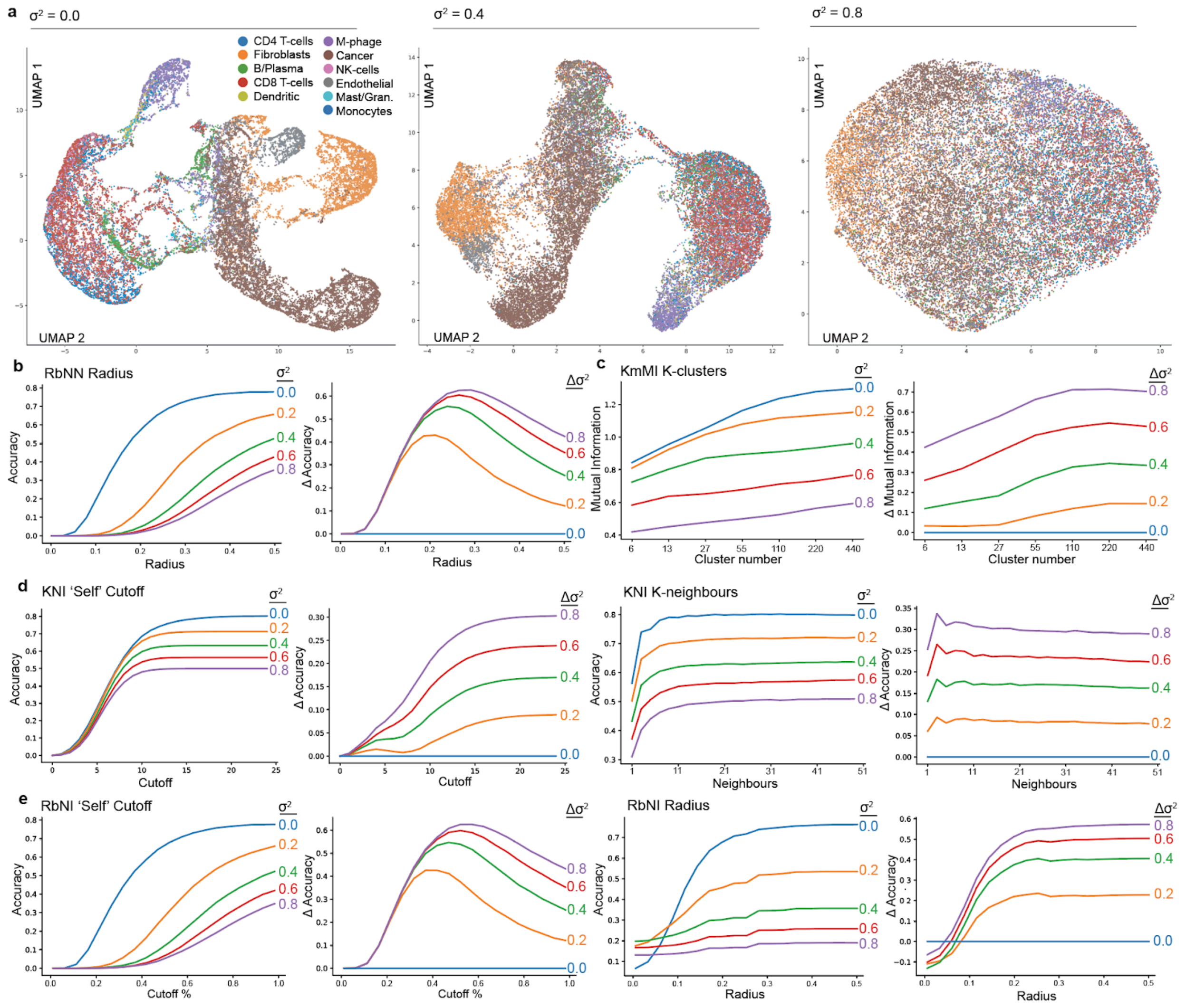

**Figure S2: Comparison of metrics on simulated noise added to a real cell-type embedding space:** a) UMAP projections of the test cases corresponding to addition of noise to the 5 dimensional embedding space, $\sigma^2 = \{0, 0.4, 0.8\}$, default UMAP parameters are used as per (McInnes, Healy, and Melville 2018), cells are coloured by 'ground-truth' cell-type; b) Batch Adjust Silhouette Score applied to the 5 test cases $\sigma^2 = \{0, 0.2, 0.4, 0.6, 0.8\}$; c) RbNN score applied to the 5 test cases (left), difference between the ideal case $\sigma^2 = 0$ and the 5 test cases, all while varying the radius parameter *r*; d) Same as (c) for the KmMI score varying cluster number; e, f) Same as (c, d) but considering the RbNI andKNI scores, where both the cutoff parameters and neighbor search parameters are varied.

***Batch effects:*** We next compared metric performance on introduction of spiked batch effects into the embedding space. To simulate progressively worse batch effects, we split the 5 dimensionally embedded data into 5 equally sized groups $C_i$, $i = \{1...5\}$, $C \in R^5$ and add a constant value μ to the respective dimension for each group where $\mu = \{0, 0.1, 0.2, 0.3, 0.4\}$ (Figure S3 a). Data was then re-normalized to unit mean and variance after the addition. The effect of this can be seen in UMAP projections of the embedding space for $\mu = \{0.0, 0.2, 0.4\}$.

1. ***Batch Adjusted Silhouette Score:*** The Silhouette score again linearly decreased from -0.0267 through to 0.0303, highlighting the silhouette score's ability to capture batch effect, however changes in the score were very small vs. in the previous example, indicating a lack of invariability to dimensionality.

2. ***RbNN:*** The RbNN method shows sensitivity to small batch-effects, however, the range of parameters over which this can be seen is much smaller (Figure S3b). Of note, in this example as the radius parameter increases beyond small values, e.g. $r = 0.15$, the score difference between $\mu = 0.0$ and $\mu = 0.1$ not only decreases but even returns to zero. At these radiuses, $\mu = 0.3$ and $\mu = 0.4$ are difficult to separate. Overall, this indicates a high degree of sensitivity to the parameter $r$ in sensing batch effects, creating a significant challenge to using this metric for assessing cell-type space alignments on real data.

3. ***KmMI:*** In this example, the KmMI metric showed even greater insensitivity to the addition of small batch effects, than it did to noise (Figure S3c). Here, greater than $\sim k > 440$ clusters were needed to even begin to see separation between $\mu = 0.0$ and $\mu = 0.1$, at this number of clusters only 70 cells on average are present per cluster. This is a number of cells to those analyzed in the local neighborhood by the Radius-based and K-neighbors metrics. This analysis indicates the KmMI as a potentially poor metric for identifying batch effects and suggests more local analysis of cell-type space approaches may be better.

4. ***KNI:*** In this example the KNI emerges as being very effective in identifying and quantifying the addition of batch effects to the aligned cell-type space (Figure S3d). Varying the cutoff number τ shows that all batch effects can be well separated for values of τ>17 (k set at 25), however, peak separation under the addition of small batch effects occurs lower, at ~τ = 12. Setting τ = 75% of the number of neighbors k, demonstrates that for k>5 there is good separation between all batch effect scenarios, again the best separation is seen for similar cutoff values in this data as is seen in the simple theoretical case given in the first section, indicating limited sensitivity in the cut-off value as the dimensionality and complexity of the data increases. Overall the KNI emerges as a good score for detecting batch effects.

5. ***RbNI:*** The RbNI also emerged as being sensitive to the addition of batch effects (Figure S3e). Similar to the KNI, sensitivity to batch effects was stable over a wide range of cutoffs percentages, $\tau^* > 60\%$ when the radius was fixed at a value of 0.3. Unlike the RbNN we also find that the RbNI is stable over a much wider range of radiuses $r$. In this example, for $0.5 > r > 0.3$ the RbNI is able to separate between all of the batch effect scenarios effectively. This indicates that the RbNI is likely a better metric for distinguishing batch effects than the RbNN.

Overall, this analysis highlights that value in KNI and RbNI approaches for assessing batch effect removal, and highlights limitations in the RbNN and KmMI methods. The silhouette score still demonstrates potential value, but is likely highly dependent on the dimensionality and complexity of the cell-type space being evaluated.

***Rescue Cells:*** In the final test, we looked at a failure case we identified in the RbNN metric. Specifically, we find that a few number of correctly labeled cells overlapping with other batches, can lead to the entire cluster being given a correct, non-outlier label, despite the presence of obvious and large batch effects. We simulate this situation by taking the largest batch effect from above, where $\mu = 0.4$ and then assigning either 1% or 30% percent of cells a different batch-label, such that clusters driven by batch-effects contain either 1% or 30% cells from other batches. We then consider how the metrics score the cell-type alignments after this mixing (Figure S4a-c).

Here we find that while the RbNN metric is highly sensitive to the mixing of even 1% of data points, (0.095 at 0% mixing; 0.41 at 1% mixing). In contrast, neither the KNI (0.016 at 0% mixing; 0.024 at 1% mixing) or RbNI are sensitive to a small number of 'rescue cells' (0.012 at 0% mixing; 0.020 at 1% mixing). This is because the number of cells in the neighborhood need to surpass the thresholds $\tau/\tau^*$ to not be given an outlier 'batch-effect' label. All three metrics assign a higher score to the 30% mixing threshold, as this represents a better aligned cell-type space.

##### *Construction of the scMARK benchmark dataset*

Prior to scREF we developed an initial smaller benchmark, that we term scMARK. We used scMARK to optimize model performance and further test the KNI and RbNI metrics, prior to our evaluation of methods on the full scREF benchmark. We also used this benchmark to initially test a wider range of model's ability to align scRNA data, including those we found less able to scale to dozens of datasets due to computational demand, should any of these have results that would justify efforts to scale them. To develop scMARK we standardized original author cell-type labels and gene identifiers across studies, such that good alignments reflected consensus labels. In total, we collected data from 11 high-quality scRNA publications from different labs, taking a 10,000 random cell sample from each study (Table S1). We recorded 29 standardized cell-type labels occurring in two or more studies and 13,865 genes in common across the 11 studies (Supplementary Methods, Table S1). 10 studies were produced using 10X Chromium technology, while (Azizi et al. 2018) was generated using the inDrop. We selected the 11 studies such that each organ, tissue and disease-type combination (eg: breast cancer/normal, colorectal cancer/normal, etc.) appears in at least two studies, and all cell-types are also present in at least two studies (Table S1). This enables us to compare alignment of every cell-type to at least one other different study. We consider scMARK an easy-to-use, benchmark dataset for testing and optimizing scRNA atlas dataset alignment methods.

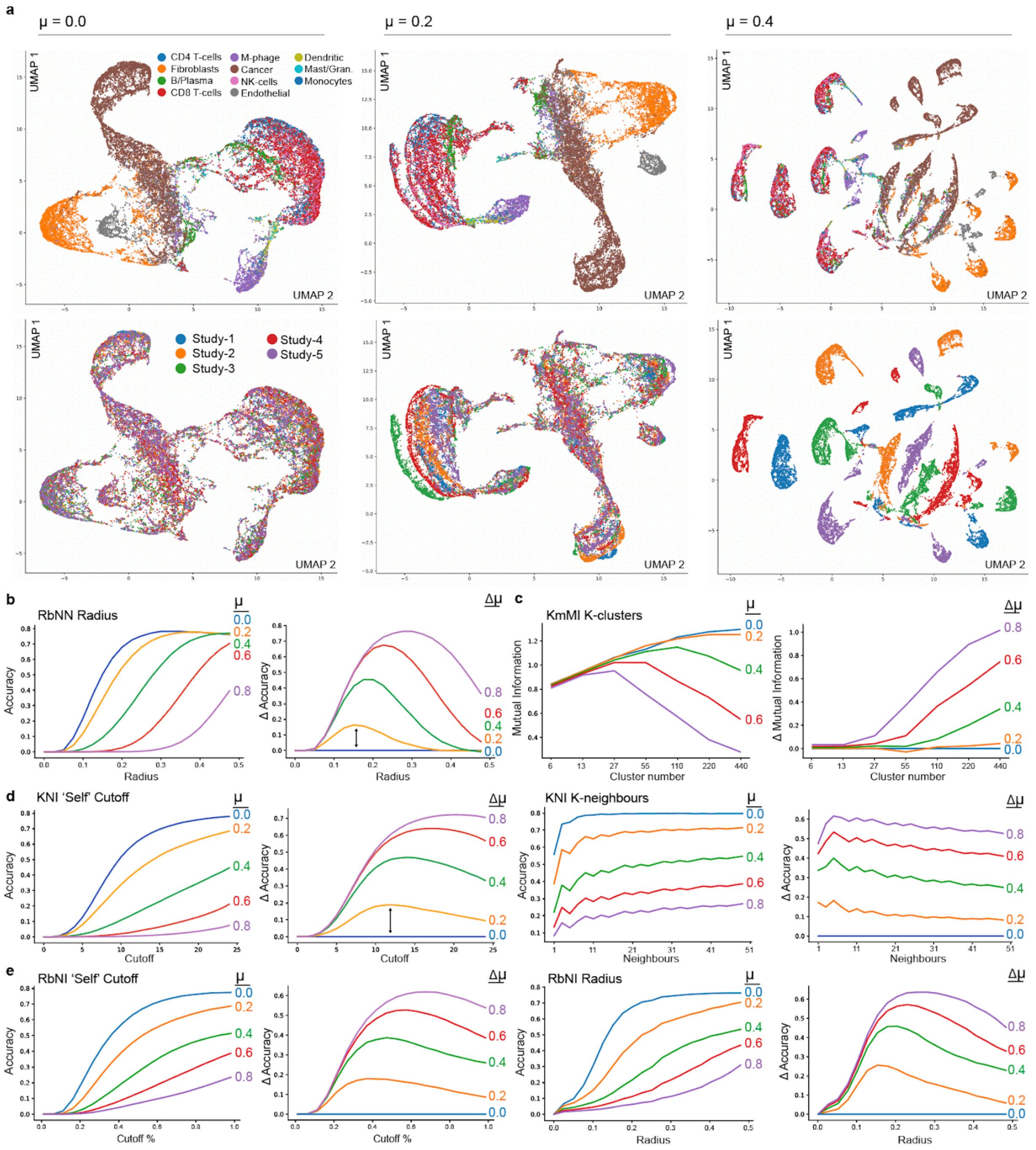

**Figure S3: Comparison of metrics on simulated batch effects added to a real cell-type embedding space:** a) UMAP projections of the test cases corresponding to addition of noise to the 5 dimensional embedding space, $\mu = \{0.0, 0.2, 0.4\}$, default UMAP parameters are used as per (McInnes, Healy, and Melville 2018), cells are coloured by simulated batch effect top panels or by coloring by ground-truth cell-type bottom panels; b) RbNN score applied to the 5 test cases (left), difference between the ideal case $\mu = 0$ and the 5 test cases, all while varying the radius parameter $r$; c) Same as (c) for the KmMI score varying cluster number; d, e) Same as (b, c) but considering the RbNI andKNI scores, where both the cutoff parameters and neighbor search parameters are varied.

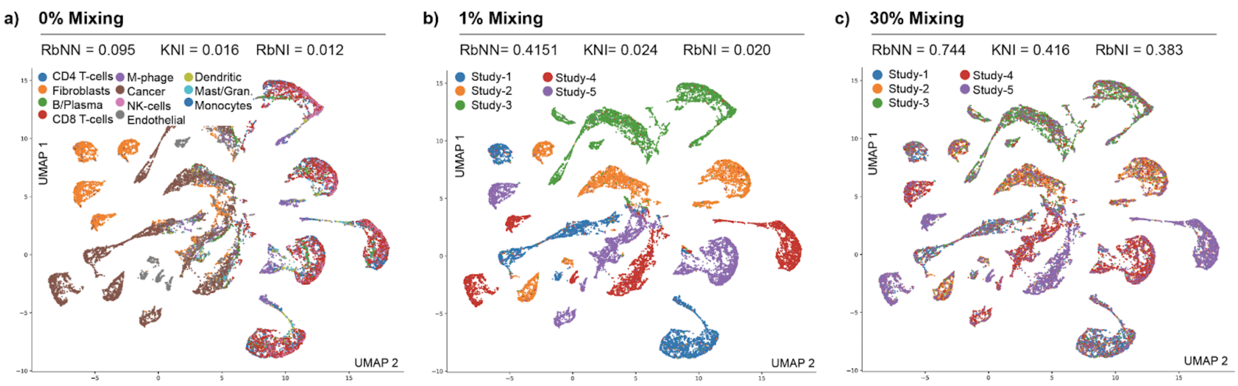

**Figure S4: The effect of swapping the batch label of a small number of cells in the presence of large batch effects** $\mu = 0.4$, here termed "rescue cells": a) UMAP projection of the embedding space with simulated batch effects and 0% mixing of cell labels, data points colored by ground-truth cell-type label; RbNN, KNI, and RbNI scores are given for this embedding space above; b) UMAP projection of the embedding space, coloured by simulated batch ID, 1% of cell batch IDs are swapped, with metric scores assigned to this space provided above; c) Same as (b) with 30% of cell batch IDs swapped.

##### *Method performance evaluation and optimization on scMARK*

Using the KNI and RbNI metrics, we initially evaluated the ability of the most commonly used methods to align / analyze scRNA data on scMARK. Namely: (1) Principal Component Analysis (PCA) applied to highly variable genes (PCA; (Kiselev, Yiu, and Hemberg 2018); (2) Reciprocal PCA as described in the Seurat workflow (RPCA; (Hao et al. 2021); (3) Scanorama (Hie, Bryson, and Berger 2019); (4) Variational Auto-Encoder with Mean Squared Error Loss (VAE MSE; included as a base-case for more advanced deep-learning tools); (5) Base-line Single-cell Variational Inference (scVI; (Lopez et al. 2018); (6) Harmony, (Korsunsky et al. 2019); and (7) An optimized scVI, where we tuned the scVI architecture against the KNI and RbNI scores. We applied these unsupervised methods to normalized gene expression matrices from the complete set of 11 scMARK studies to generate a unified cell-type space. We then used the RbNI and KNI metrics to evaluate the quality of this cell-type embedding space.

Based on this framework and selection of the top performing model after varying key parameters, we found that an Optimized scVI model and Harmony outperform other methods at aligning the scMARK dataset under both the KNI and RbNI scores (Figure S5a; Table S2). Qualitatively, in UMAP projections of the cell-type embedding space, scVI and Harmony were the only models where different cell-types formed separate clusters and where datasets did not form their own specific groupings, indicative of batch effects (Figure S5c, d). We also tested alternative embeddings dimensionalities for RPCA, Harmony, and Scanorama in case results improved noticeably here, but found this was not the case (Table S2; KNI only since the RbNI is sensitive to dimensionality). Thus, we show the value of both the Harmony and scVI cell-type methods for alignment and the RbNI and KNI metrics for measuring alignment quality. Model details and parameters are given in Methods.

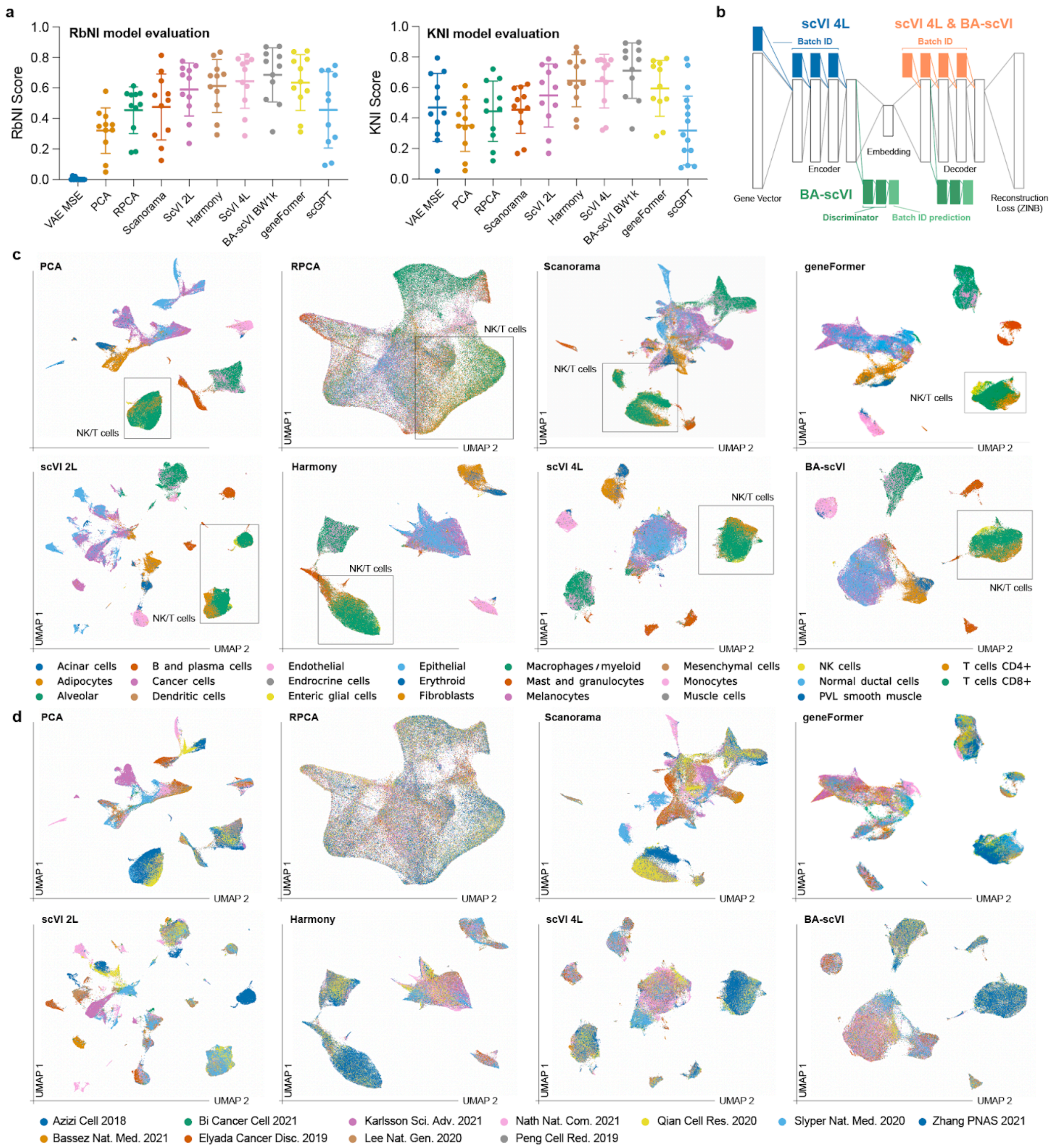

**Figure S5. Comparison of Model Performance on scMARK:** a) The K and Radius-based Neighbor Intersection (KNI and RbNI) scores were used to assess scRNA alignment model performance on scMARK's 11 high-quality scRNA datasets. KNI and RbNI scores are plotted for each of the 11 datasets, alongside the mean score and standard deviation, for each alignment method. A perfect score is 1; b) To improve the performance of scVI under the KNI and RbNI metrics, we introduced BA-scVI in which discriminators are trained to identify batch effects in layers before and after the cell-type embedding layer (green) vs. scVI where batch ID vectors are injected into encoder and decoder layers (blue); c) UMAP projections of alignments produced by the eight different methods where cells are colored by author provided 'ground-truth' cell-type labels; d) UMAP projections as in ©, but colored by study. The NK/T-cell grouping is highlighted to show variation in cell-type alignment quality between the methods.

##### *RbNI / KNI breakdown into Batch and Accuracy metrics highlights Harmony and scVI limitations*

In the UMAP projections of the scVI alignment (Figure S5c, d ) we noted significant batch effects still existed in more challenging datasets such as Azizi 2018 (generated using InDrop). Meanwhile, in the harmony alignment, we observed B-cells were seen to overlap with the larger T-cell grouping, indicating overfitting. Indeed, when we broke the KNI and RbNI scores down into the respective kBET and accuracy components, we found that while both the 2L and 4L variants of scVI performed better than Harmony on accuracy, Harmony outperformed at batch-effect correction (Table S2; Figure S6 a-f). This confirmed our qualitative observation, and further supported the inclusion of direct batch penalization in the scVI model via an Adversarial approach via BA-scVI.

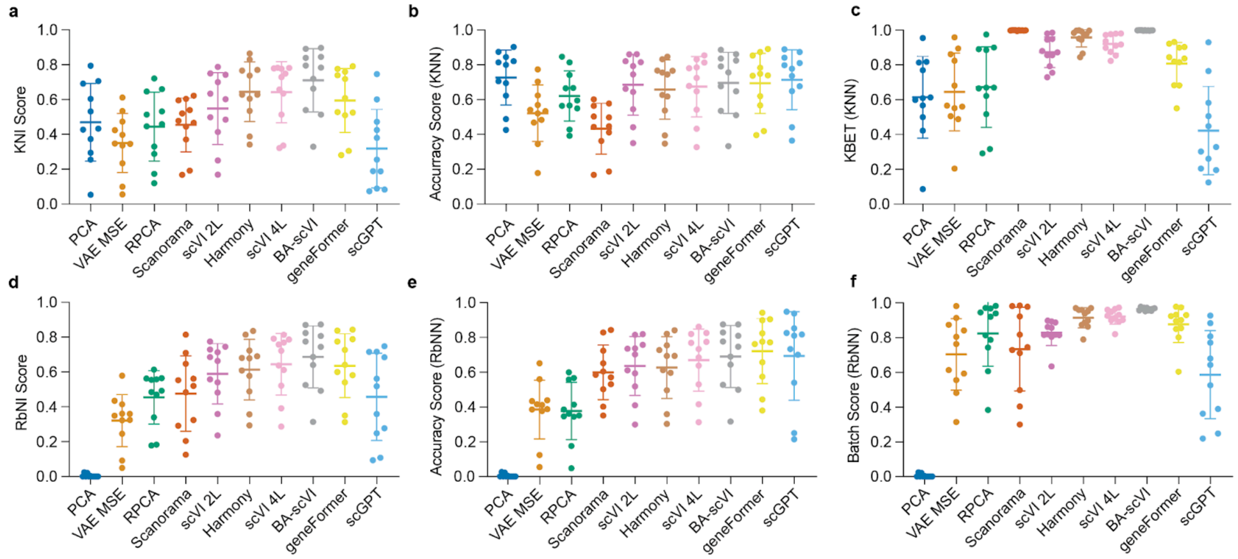

**Figure S6: Comparison of metrics on cell-type embedding spaces generated by models evaluated in the main text, broken down by accuracy and batch-effect correction** a) KNI scores as shown in Figure S5a; b) Accuracy score only, calculated from prediction of held-out cell-type by K-nearest neighbors (Methods); c) Batch correction only, calculated as the kBET score (Büttner et al. 2019); d) RbNI scores as shown in as shown in Figure S5a; e) Accuracy score only using a radius based nearest neighbors; f) Accuracy only using radius based nearest neighbors to calculate the kBET score.

##### *BA-scVI optimization*

The above studies led us to adopt an adversarial training regime for scVI described in Methods and the main text. scMARK was used to optimize the BA-scVI architecture prior to evaluating it on scREF. Specifically we identified an optimal BA-scVI architecture for aligning scMARK, consisting of 4 hidden layers in the encoder and decoder networks, and 2 layers in the adversarial batch-detection network (Figure 1b, BA-scVI 4L Both; Table S2, Figure S5b). Importantly, on scMARK, BA-scVI significantly outperformed an optimized scVI architecture (RbNI, P=0.005; KNI, P=0.012; paired T-test) that also consisted of a 4-layer encoder and decoder (Table S2). This result was consistent over different learning rates and stopping criteria under both the KNI and RbNI scores (Table S2). On assessment of the BA-scVI accuracy and KBET score, we found that BA-scVI both outperformed scVI on alignment, and Harmony on batch effect correction, thus giving further confidence in this approach to alignment at scale (Figure S6b, c, e, f). We also consistently observed that the number of layers, choice of batch identifier, and weighting on the batch penalty were key determinants of the KNI and RbNI scores obtained (Table S2). Qualitatively, in UMAP projections BA-scVI resolves cell subtypes, e.g., CD4+ vs. CD8+ T-cells on difficult datasets e.g. Azizi 2018, while removing any clear batch groupings (Figure S5d). We performed ablation studies; and confirmed the application of adversarial constraints to the encoder and decoder performs best (Table S2). We also tested batch cost functions and weights and found that cross-entropy loss performed best (Table S2). Together this optimization results in the BA-scVI architecture we use to align the benchmark datasets (Figure 1b; Main Text).

##### *Single-cell transformer model testing*

In the course of this work, two transformer models were released, namely geneFormer (Theodoris et al. 2023) and scGPT (Cui et al. 2024). We evaluated: (a) The high-dimensional embeddings; (b) The author recommended an approach to fine-tuning for batch-effects (Methods); and (c) a BA-scVI model trained on the embeddings. In all cases BA-scVI performed better than these large-language model based approaches on scMARK (Table S2; Figure S5). The number of epochs and training time required to converge also increased following embedding of the raw values with geneFormer and scGPT, while the KNI and RbNI scores decreased, potentially driven by the injection of batch-effects that could not be removed by the BA-scVI model applied to the transformed embedding space. This suggests at best these transformer models have limited value in aligning scRNA at scale (Table S2), and may in fact be further compounding the problem of batch effect removal / deconvolution. We note we did not perform a thorough evaluation of alternative potential reconstruction loss options for training BA-scVI on the transformer generated embedding spaces, and only used cosine similarity as per fine-tuning procedures in the original studies (Theodoris et al. 2023; Cui et al. 2024), this may represent a route to improving results. In the main text for scREF evaluation we use the author provided approach to fine-tuning coupled to an autoencoder for dimensionality reduction to evaluate scGPT, and scVI (3-layer) trained on the geneFormer embedding space to fairly test these embedding spaces for scRNA alignment tasks.

##### *Analysis of metric correlations on the scMARK Benchmark Dataset*

As a final assessment of metric reliability, we looked at how well metric scores correlated with each other across the set of models tested on the benchmark dataset, and considered cases where metric scores did not correlate well. Specifically, we compared scores obtained by the set of metrics (KNI, RbNI, Silhouette Score, KmMI, and RbNN) over all deep-learning architectures tested, with learning rate = 1E-4, and patience = 100, as well as RPCA and PCA methods (Figure S7a; see methods for parameters).

- ***RbNI, KNI and KmMI Scores:*** We found that the RbNI and KNI scores correlated well with each other, this was largely expected due to the similar way in which both scores are calculated. The KmMI score also correlated highly with the RbNI and KNI scores; this providing support to the notion that the RbNI and KNI provide robust measures of cell-type alignment quality (Figure S7a).
- ***Batch Adjusted Silhouette Score:*** In contrast to the RbNI, KNI and KmMI score, the Silhouette score showed negative correlation with all other metrics. To better understand this we explored the UMAP space of a method that performed highly under the Silhouette score but poorly under the other three (ScVI 2L, No

batch ID, No library encoding). A UMAP projection of this space indicated significant batch effects still existed in this cell-type space and alignment of author labels was poor. It is unclear why the silhouette score assign a high value to this cell-type space, but under visual inspection with UMAP, projections generated by the top performing model under the RbNI, KNI, and KmMI (BA-scVI 4L, with 2 hot vector encoding 'Both' sample ID and study ID), generated much better alignments. *Thus we propose the Silhouette score is a poor metric by which to evaluate cell-type embedding spaces.*

- ***RbNN Score:*** The correlation between the RbNN score and the other three metrics was higher, but clear differences existed. We examined the case of the cell-type space generated by BA-scVI 2L, with 2 hot vector encoding 'Both' sample ID and study ID), that performs well under RbNN evaluation (score = 0.7214, rank = 3), but poorly under the other three metrics (KNI, score = 0.7214, rank = 22; RbNI, score = 0.5677, rank = 16; KmMI, score = 0.8067, rank = 23). An assessment of the cell-type embedding space here with UMAP indicates that significant batch-effects still exist, especially in the Azizi study performed on the InDrop. However, excluding this study, a number of datapoints clearly still overlap with the Azizi dataset and are correctly labeled. Overlaying an illustrative mask that depicts a radius around the overlapping data points demonstrates how these sparse overlapping data points may 'rescue' the overall RbNN score as shown in the *Rescue Cell* the example above (Figure S7b. Thus, the KNI, RbNI, and KmMI scores provide key advantages over the RbNN score.

##### *The RbNI and KNI Scores emerge as intuitive and effective metrics for evaluating cell-type spaces*

Across the metrics tested, the RbNI and KNI emerged as powerful metrics that fulfill all the requirements needed to evaluate the quality of an aligned cell-type space as defined here. An initial study of the theoretical case of two cell-types and two batch effects demonstrated the need for scnRNA alignment evaluation metrics to explicitly penalize batch-effects. The Batch Adjusted Silhouette Score, KmMI RbNN, RbNI and KNI were then all able to effectively separate batch effects in the simple example considering two batch-effects, and two cell-types. In the more realistic case, noise was spiked into a single-embedded scRNA dataset, to simulate less accurate grouping of cell-type all of these methods also were able to detect poorer cell-type alignment. However, in the case of detecting batch, the RbNN methods demonstrated an acute sensitivity to selection of the parameter $r$ while the KmMI score required a very high cluster number to identify the addition of small batch effects. The RbNN score also was sensitive to the ability of a small number of cells to 'rescue' large batch effects. Thus, in these theoretical and simulated examples, the RbNI and KNI emerged as the best metrics for scoring the alignment of scRNA datasets. The silhouette score also emerges as a potential score of alignment quality, though was clearly highly sensitive to dimensionality and dataset complexity.

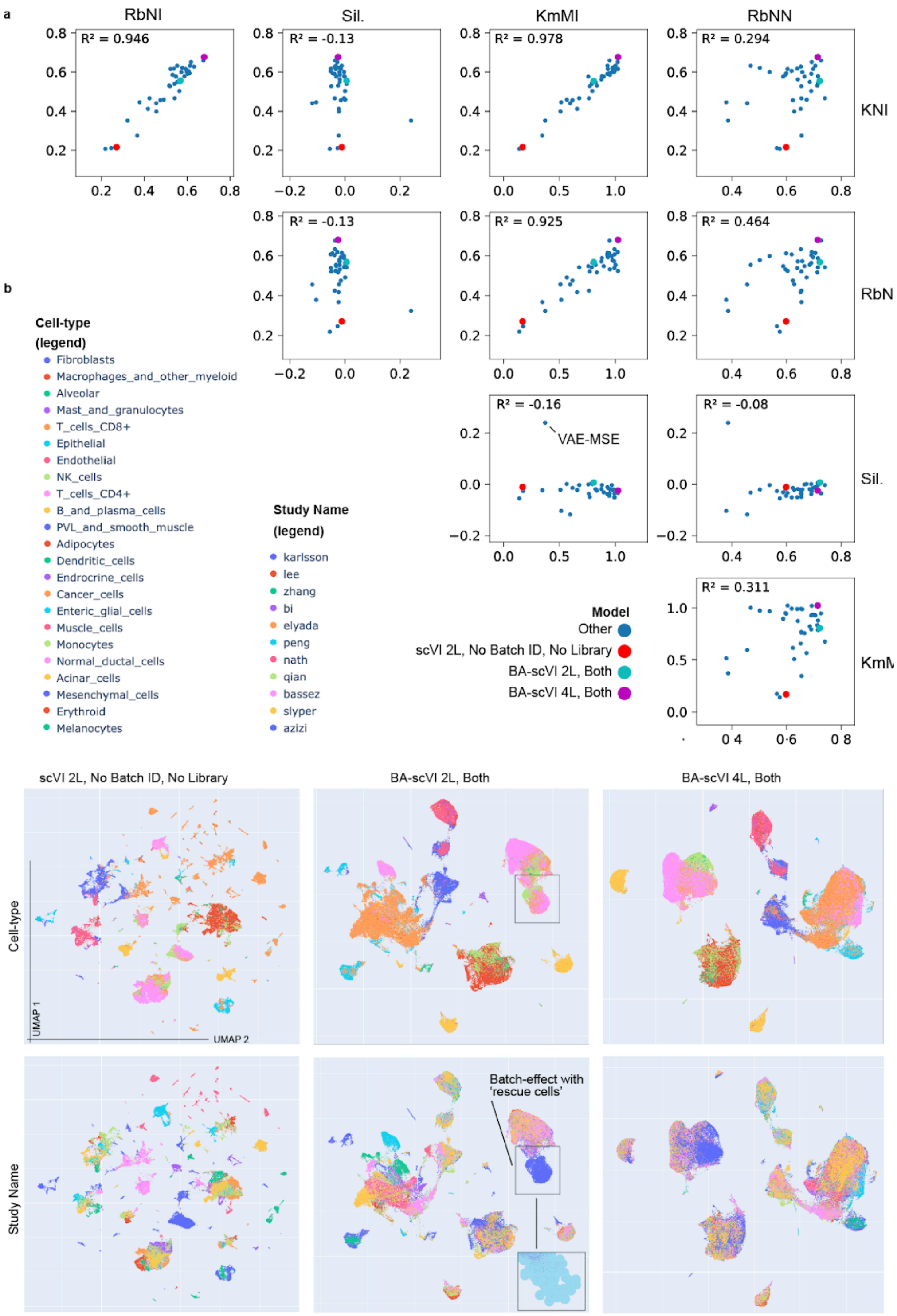

**Figure S7: Comparison of metrics on cell-type embedding spaces generated by models evaluated in the main text, and deep-learning architectures considered during optimization of the scVI and BA-scVI models:** a) Correlation scatter plots between scores generated by the different metrics; example cases plotted in (b) are coloured uniquely; b) UMAP projections of example cases in (a) coloured by ground-truth cell-type label (top row), and study name (bottom row), a case of rescue cells leading to a high RbNN score despite the presence of large batch effects is illustrated in BA-scVI 2L Both, by removing the Azizi dataset (dark blue) and plotting a fixed radius circle (nominal) around remaining 'rescue-cells'.

In our evaluation of these metrics on the alignments generated by models on our benchmark dataset we noted that the RbNI, KNI, and KmMI scores are highly correlated, while the Batch Adjusted Silhouette score and RbNN score generate very different results. A comparison of UMAP projections of the aligned cell-type spaces indicates that the RbNI, KNI and KmMI scores are much better at identifying cell-type alignments that result in interpretable outputs. We note this analysis may be biased toward metrics that align well with the UMAP projection technique. Indeed, the RbNI calculates alignment quality in a manner similar to that by which UMAP reduces dimensionality, i.e., using a fuzzy-radius based neighbors approach (McInnes, Healy, and Melville 2018). Given UMAP and the closely related projection method tSNE are the dominant means of projecting and visualizing scRNA data in use today, we feel metrics and alignment models that perform well under UMAP are most valuable to the community, though note re-evaluation of the models and metrics we present here is warranted if other visualization tools become similarly established. Thus from this analysis and the example cases before the RbNI, KNI and KmMI emerge as the best metrics for scRNA alignment evaluation.

The RbNI, and KNI make intuitive sense and can be used to guide scRNA alignment model development. Specifically, the reasons that a datapoint is labeled incorrectly can be identified and in turn optimized for. Namely:

1) ***KNI and RbNI:*** Inaccurate cell-type labeling, e.g., cases where a CD8+ T-cell is labeled as a CD4+ T-cell. This failure is driven by either poor-cell-type space alignment or over-alignment such that different cell-type clusters overlap. Here, better model accuracy is needed.

2) ***KNI and RbNI:*** Cell-type is labeled as an outlier due to having too many cells from the same dataset nearby. This failure is driven by the presence of batch-effects, here improved alignment / batch-effect correction is needed.

3) ***RbNI only:*** Cell-type is labeled as an outlier due to being too far from other cell-types. This failure is driven by loose clustering, and indicates poor global alignment, i.e., clusters are dispersed, vs. tight and specific to cell-types. Models need to be developed that generate tighter clustering within cell-types vs. between cell-types (normalization of the data prior to evaluation ensures distances are relative vs. absolute).

Additionally, with the KNI and RbNI metrics a score per-dataset can easily be output, vs. the global scores given by Silhouette Score, and KmMI, breakdown can also be performed on a per cell-type basis as we use to asses FOXP3+ T-cell labeling (Figure S10), again another useful advantage. Overall, we thus find the RbNI and KNI as the most valuable metrics for testing the quality of aligned cell-type spaces based on this work. We therefore focus on the RbNI and KNI scores for evaluation and testing of models in the main text on the scREF dataset.

Given the top performing model (BA-scVI 4L Both) is top under all three metrics, vs. discrepancies emerging, and the fact that this score is difficult to interpret vs the RbNI and KNI, attention to the KmMI is not given in the main text. We also note that the Adjusted Rand Index (ARI) is often used to evaluate scRNA alignment quality, and could be substituted for Mutual Information in calculating this score. Given the KmMI score generates valuable results, and substituting in the ARI for MI would not lead to an improvement in interpretability, we feel an assessment of ARI vs. MI would not lead to a meaningfully more valuable metric and is also not warranted here.

##### *scVI and BA-scVI qualitatively result in the best alignments of scREF as assessed by UMAP*

Calculation of the KNI and RbNI scores of scREF alignment by the 6 models, BA-scVI, scVI, PCA on Highly Variable Genes (HVG), Harmony applied to PCA (HVG), as well as geneFromer and scGPT with BA-scVI applied to the transformed genes, found quantitatively that scVI and BA-scVI perform best at aligning scRNA data at scale (Table S3). Here we find qualitatively that cell-type spaces generated by scVI and BA-scVI show a high-level of separation between cell-types, as well as overlap between studies, indicating successful alignment (Figure S8). Of note, we see a small but noticeable improvement in overlap in BA-scVI alignments vs. scVI; e.g. in myeloid cell-types, (Figure S8). In contrast, using the transformer models scGPT and geneFormer to embed the cell-type space prior to fine-tuning as per author protocols, resulted in noticeably worse alignments where batch effects could clearly be seen (Figure S8), BA-scVI training on the embeddings space did not improve results (Table S3). Calculation of the KNI on the raw fine-tuned embedding space also showed poor results (Table S3), further highlighting the limitations of these methods. UMAP applied to PCA (HVG), led to a surprisingly good alignment at scale, where major cell-type groupings could be resolved, however batch effects were present. Harmony corrected very well for batch effects, but cell-types could not be clearly distinguished. Thus, qualitatively we also find that scVI and BA-scVI are most effective at aligning scRNA atlases.

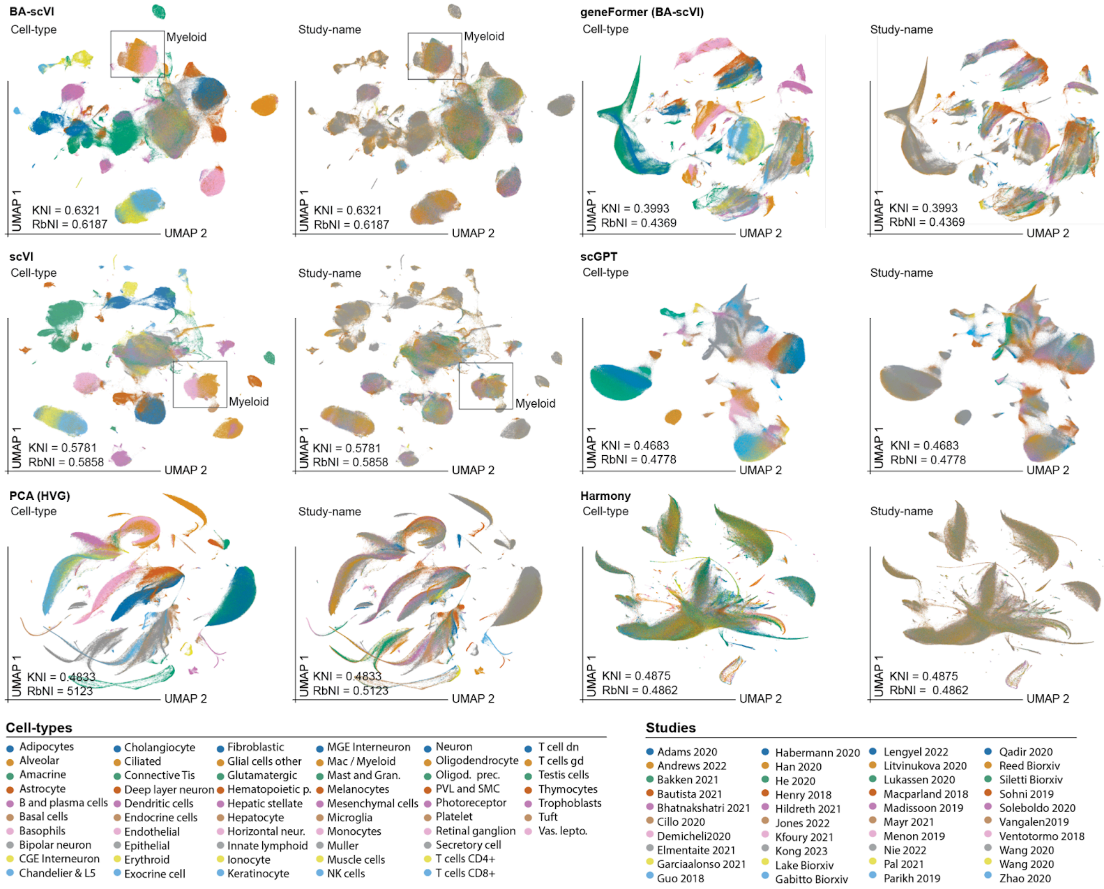

**Figure S8: UMAP projections of scREF embedding spaces for the set models presented in Figure 1:** UMAP projections of alignments produced by the 6 different methods compared in the main text, in the left panel for each method, cells are colored by standardized author provided 'ground-truth' cell-type labels, in the right pane cells are colored by study.

##### *Rare cell-type groupings emerge in cell-types spaces aligned by BA-scVI*

In characterizing the cell-type embedding space generated by BA-scVI early on we sought to understand whether outlier groupings could be explained by biological variation or were due to technical artifacts. To determine this we manually assessed expression of marker genes on the UMAP plot, leveraging known marker gene-signatures in the human protein atlas (Karlsson et al. 2021). Here we identified three genes that correlated with outlier groupings, namely, CLEC4C, MPZ, and CRYBA2 (Figure S9a, b). Human Protein Atlas labellings (Karlsson et al. 2021) and literature review determined these cells were likely: 1) Plasmacytoid Dendritic Cells (Dzionek et al. 2001); 2) Schwann cells (Lemke, Lamar, and Patterson 1988); and 3) Colon Endocrine cells (Beumer et al. 2020) respectively. To further validate this finding, the expression pattern of highly correlated genes based on the Human Protein Atlas scRNA dataset (Karlsson et al. 2021) was then also assessed in the cell-type embedding space. Here we identified CLEC4C+ cells, as also expressing high levels of LILRA4, and PTCRA, known markers of Plasmacytoid Dendritic cells (Murray, Xi, and Upham 2019; Dzionek et al. 2001; Shigematsu et al. 2004), giving us confidence in this cell-type labeling (Figure S9c). Examining the MPZ+ cell-type grouping we determined that these also express high-levels of the markers FOXD3 and SOX10, also consistent with this cell-grouping being Schwann cells (Furlan and Adameyko 2018). Finally, CRYBA2+ cells also expressed somatostatin (SST) and CHGA known to be expressed in Colon endocrine cells at high-levels, (Kasprzak 2021; Nagatake et al. 2014), in line with this being a cell-grouping. Overall, this analysis provides evidence these cell-type groupings are indeed rare cell-type vs. technical artifacts, and defines the markers we use to validate rare-cell types can be seen in the aligned scREF atlas.

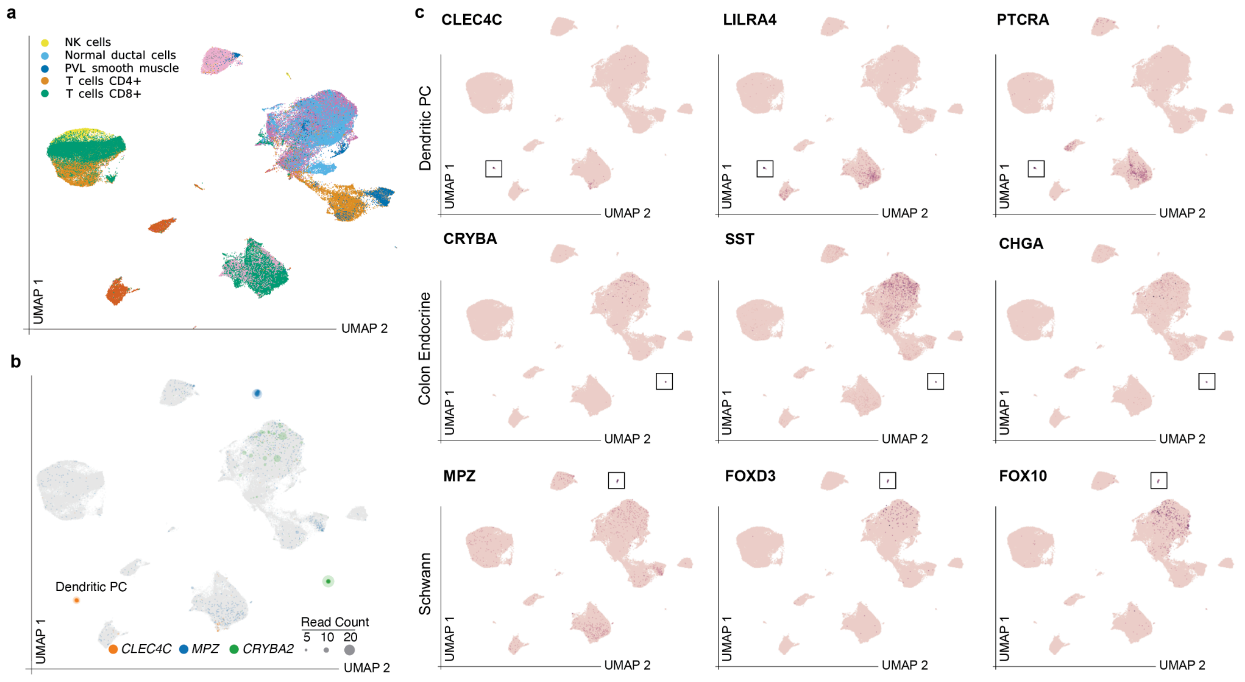

**Figure S9: Expression of rare cell-type marker genes:** UMAP projections of the cell-type embedding space coloured by log gene expression of rare-cell type marker genes. Outlier groupings were seen in the BA-scVI embedding space. An analysis of known rare-cell type marker genes identified three ‘anchor’ genes that uniquely localized to each of these outlier groupings; this level of consistency in marker expression indicates these cell-type groupings correspond to rare-cell types vs. technical artifacts.

##### *BA-scVI performs well at the discovery of cell-types not in the set of standardized cell-type labels*

A key assumption in our benchmarking approach is that consensus author labels can be used to identify models that align scRNA data effectively, and that the KNI and RbNI readouts capture this. FOXP3+ T-cells are a well-characterized CD4+ T-cell subtype, termed regulatory T-cells (T-regs), that are not defined in scMARK benchmark dataset standardized cell-type label set. Following cell-type alignment with BA-scVI, we found that within the CD4+ T-cell cluster, FOXP3 expressing cells localize to a distinct region (Figure 10a, b), indicating that cell-types outside of the standardized set can be discovered in the unified space. Quantitatively, if we set FOXP3+ T-cells as a new cell-type based on a threshold gene expression score, we see that BA-scVI also outperforms other methods at identifying this additional, cell-type based on the KNI and RbNI scores (Figure S10c; Table S3). Overall this indicates that consensus cell-type labels can act as a good readout for the alignment quality of ‘new’ cell-types, thus supporting both the initial assumption that they are a good ground truth and the value of the KNI and RbNI metrics for identifying models that perform well in a discovery setting.

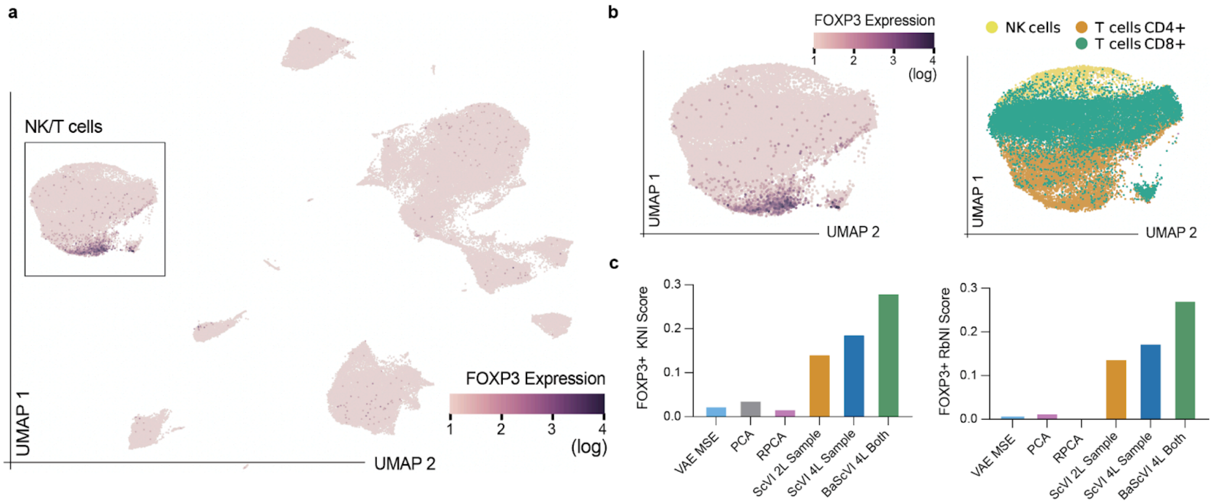

**Figure S10 Analysis of BA-scVI on cell-type outside of the standardized set and on cell-types with a known ground-truth (*in vitro* data):** ) UMAP projection of scMARK’s 11 datasets colored by expression of T-reg marker FOXP3; b) Close-up of selective localization of FOXP3+ cells to CD4+ T-cell clusters; c) Performance of alignment methods at labeling cells with FOXP3+ expression above a UMI count of 1 under the RbNI and KNI metric.

##### *Cross-species alignment enables labeling of mouse and human cerebellar granule cells*

In assessing the scREF-mu alignment we noted that a high-quality scRNA study of the mouse cerebellum (Kozareva et al. 2022) contained a number of cerebellar granule cells that failed to align well to glutamatergic neuron populations in other regions of the brain, indicating a distinct transcriptional profile for this cell-type, unique to the cerebellum. We thus excluded this cell-type from determination of the KNI and RbNI scores in the mouse and human alignments. We also noted that only a single dataset in the human atlas contained cerebellar tissue samples (Siletti et al. 2023). In the scREF alignment cerebellar cell-types are present in incredibly low abundance due to these samples accounting for only 3 out of 105 samples in the overall dataset. However, we sought to test whether upsampling of cerebellar tissue from (Siletti et al. 2023) in the scREF atlas could be used to map cerebellar granule cells from (Kozareva et al. 2022). Indeed upsampling of this tissue type and training of BA-scVI on the new joint dataset resulted in effective alignment of mouse cerebellar cells in the mouse dataset to upper rhombic lip neurons in the human dataset (Figure S11a-c). Quantitatively, we found that we obtained KNI and RbNI scores of 0.82 and 0.84 respectively on granule cells, where only cell-type labels in the other species could be used for prediction. We also found a number of cerebellar neuronal cell-types labeled as ‘Splatter Neurons" (Siletti et al. 2023) map closely to cerebellar Golgi and Purkinje cell groupings in (Kozareva et al. 2022), indicating potential mislabelling of this cell-type group in the human study. Overall, this indicates that cross-species alignments can serve to help better understand cell-types and identify consensus, especially in cases where information for specific tissue types and/or organs is less abundant.

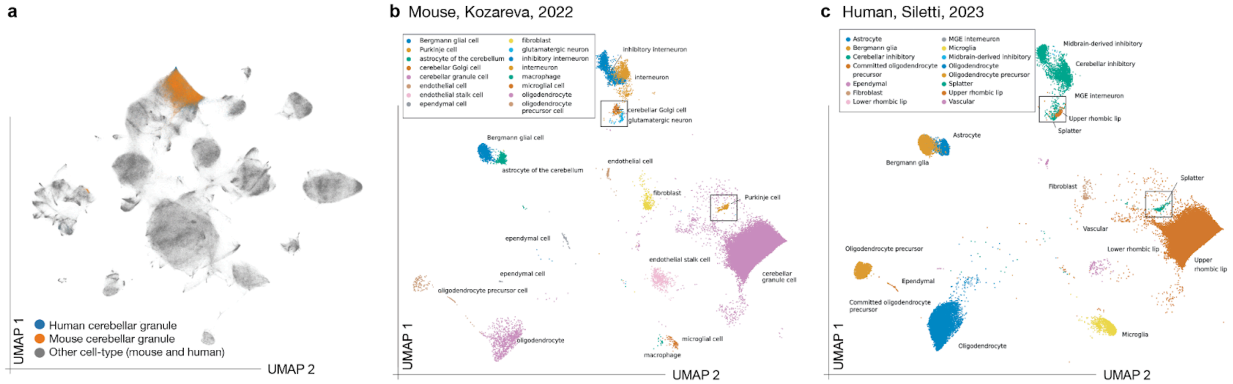

**Figure S11: Identification of a unique cerebellar granule cell population via cross-species alignment:** a) UMAP projection of BA-scVI jointly trained on scREF and scREF-mu with upsampling of cerebellar tissue in the Siletti et. al dataset. Human granule cells (upper rhombic lip; blue) and mouse cerebellar granule cell-types (orange) are labelled and quantitatively show a high-level of overlap; b) The same UMAP projection as (a) colored by author cell-type labels as per (Kozareva et al. 2022); c) The same UMAP projection as (a) colored by author cell-type labels as per (Siletti et al. 2023).

##### *Label propagation leads to clusters that closely align with author and standardized cluster labels*

To build further confidence that the number and variety of cell-types found in the aligned atlas by label propagation (LP), was not dramatically different to that of either the mouse or human atlases, we assessed LP and K-means cluster labellings against standardized and author cell-type labels for both the mouse and human atlases individually. Qualitatively we assessed UMAP projections of the four labelings for the individually aligned atlasses. Here we saw that on both the human (Figure S12a) and mouse atlases (Figure S12b), LP results visually aligned much closer to standardized labels, and author labels than K-means clustering did. Importantly in LP labellings we noted significant variance in cluster size as we would expect from analysis of tissue type containing common and rare cell-type variants, vs. K-mean clustering that resulted in excessively uniform clusters sizes.

Quantitatively, we used the Adjusted Rand Index (ARI) to measure the clustering similarity between LP and K-means, against the standardized and author labels. For both the human and mouse atlas, the ARI between LP and the standardized and original author labels was dramatically higher than that for K-means, indicating this method more accurately captures cell-type variation seen in the original unaligned datasets (Figure S12c, f). Moreover, giving further confidence in LP, we found that for the human dataset the ARI between LP and standardized labels was higher than that between standardized labels and author labels, suggesting LP derived results accurately capture biological variation, providing further confidence in LP for unbiased cell-type clustering.

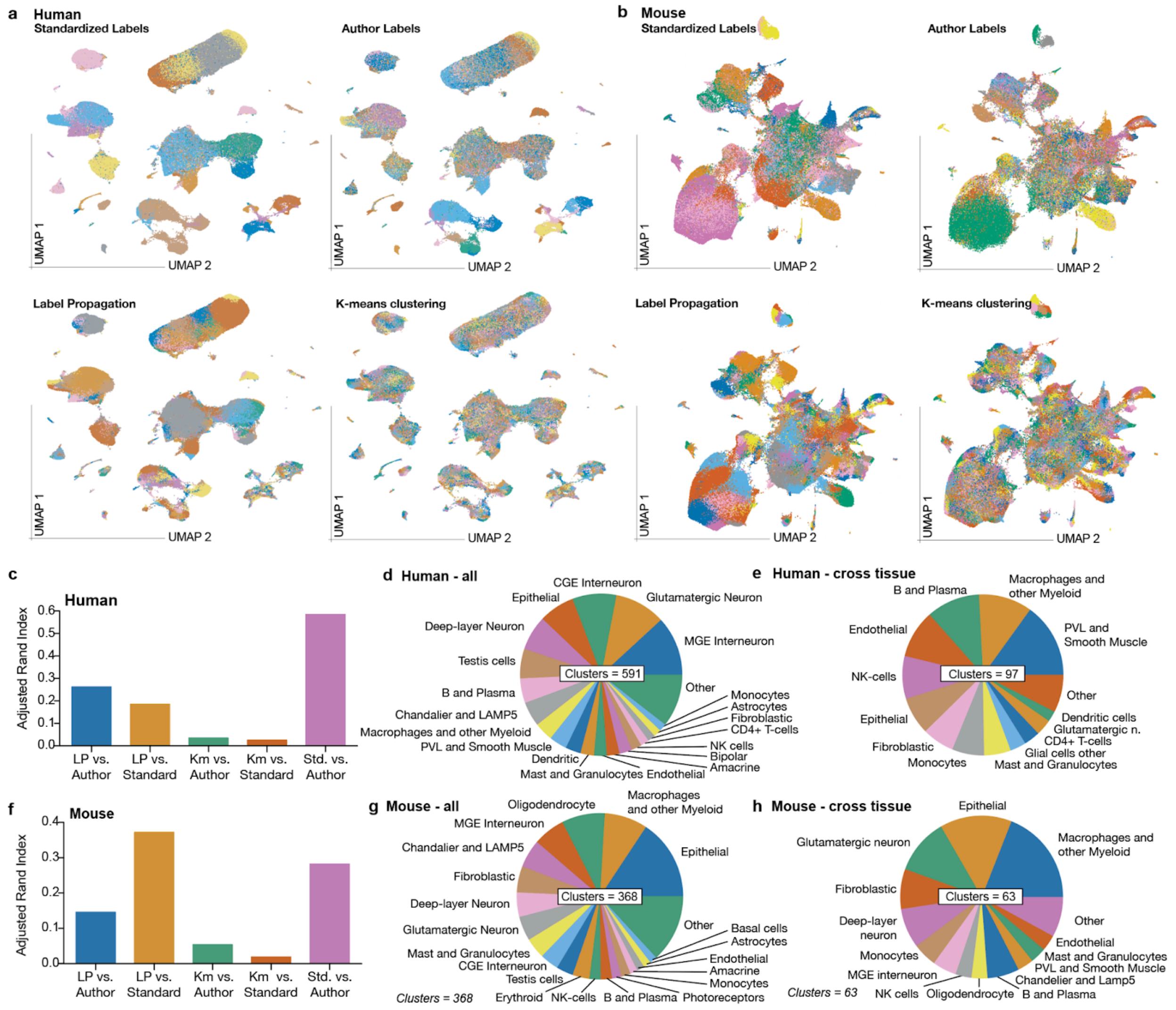

**Figure S12. Unbiased cell-type clustering of individually aligned scREF and scREF-mu atlases:** a) UMAP projections of author and standardized labels, compared to cell-type clusterings generated via label propagation and K-means clustering respectively for the human scREF atlas, ARI scores comparing the two clustering approaches with the two labellings, alongside the labellings compared to each other for reference; b) The same plots and comparisons as (a) but for the scREF-mu atlas; c) Comparison of label propagation and K-means clusters with author or standardized cell-type labels, measured by the Adjusted Rand Index for the human atlas alone; d) Proportion of unique cell-type groups, with more than 20 cells present in two datasets, by standardized label for the human atlas alone; e) Same as (d) but filtered for cell-types with over 20 cells present in 2 datasets, in more than 10 tissues in mice and humans; f-h) The same plots as (c-d) with analysis performed on the mouse scREF-mu atlas.

##### *Label propagation on scREF / scREF-mu finds similar cell-type diversity to that of the aligned atlas*

We also assessed how similar the total cluster number identified by LP was between the individually aligned moused and human atlases and the cross-aligned atlas analyzed in the main text. We determined both the distributions of cell-types emerging from LP clustered data broadly across all tissues, and filtered for those groupings present in more than 10 tissues as per the main text analysis (Methods). Across all tissues we maintained cell-type groupings with at least 30 cells (human), or 20 cells (mice) present in at least two datasets, with these different cutoffs reflecting the differences in the total dataset size for each species. In analysis of the human tissue dataset 591 clusters were found, while for mice 368 clusters were seen. In the cross aligned dataset 792 clusters were detected, and 650 of these contained at least 20 cells from a study in the mouse and 30 cells from a study in the human dataset. That the number of groups from mouse and human tissue data is on the same order of magnitude provides confidence that label propagation is reliably finding 100's and not 1000's of cells in the embedded atlases. Of note, where in the human and cross aligned atlas CNS neuronal cell-types accounted for the largest share of cell-types, this was not the case in the mouse atlas (Figure S12d, g). This is likely due to the fact that 2 large human brain atlases were curated for the scREF dataset (Gabitto et al. 2023; Siletti et al. 2023), where datasets covering the mouse brain, typically focus on specific regions, additional studies covering the mouse brian would likely increase the LP cell grouping count significantly for the mouse atlas alone.

Following further filtering for cell-types being present in at least 10 tissue types, 97 cell-types remained in the human atlas and 63 in the mouse atlas. In line with analysis of the cross-tissue atlas, epithelial cells alongside macrophage and other myeloid cell-types accounted for the most common cell-type groupings in both individually aligned atlases (Figure S12e, h). More broadly similar cell-type diversity was seen between these atlases and the cross aligned indicating consensus cell-types emerge from alignment. Of note, CNS neuronal clusters also were present in the conserved mouse tissue-types, this may be due to the lower alignment quality obtained on the mouse atlas only, as per Figure 3a, leading to cell-types misaligning with CNS tissue. We also assessed the total cell-type number represented by these conserved groupings in the mouse and human atlases. In the human atlas - conserved cell-types accounted for 2.1m of a total of 5.8m cells in the human atlas LP was run on (note we downsample the large brain human atlases to fairer balance human and mouse representation), while in the mouse atlas these conserved cell-types accounted for 800k of 3.1m cells (Table S5). Thus in both cases, these broadly conserved cell-types account for a significant percentage of all cells in the benchmark datasets.

##### ***Complement expressing myeloid and fibroblast cell-types exist in the individually aligned atlases:***

To further assess the validity of our finding that conserved complement expressing fibroblast and myeloid cell-types exist between mice and humans we also sought complement expressing myeloid and fibroblast cell-types in the scREF (human) and scREF-mu (mouse) atlases aligned individually. Here we found that indeed in both mice and humans, myeloid lineage (Labeled Myeloid cells, Monocytes, Dendritic cells) regions rich in expression of genes encoding the Complement 1Q components, C1QA, C1QB, and C1QC (Human, Figure S13a, b; Mouse, Figure S14a, b; Table S5). Inspection of gene-expression heat maps covering the major standardized cell-types across the set of tissue-types present in the human atlas (Figure S13e) further shows that this expression is largely unique to these cell-type across all tissues, although we note a low level of complete C1QA, B and C expression by PVL/SMC's in adipose tissue and expression of C1QA/B with a low level of C1QC by liver epithelial cells (distinct from liver hepatocytes); other cell-types only express a subset of these genes that may be noise due to low cell numbers. In the mouse atlas (Figure S14e), board myeloid and monocyte expression of C1QA, B, and C was seen, alongside mast and granulocytes in head and neck tissue also showed expression of the C1Q components. Thus only myeloid and monocytic cells show conserved C1QA, B, and C expression between mice and humans.

In fibroblasts we identified a conserved signature of C1R, C1S and CFD marking a large cell-subtype grouping. Specifically, UMAP projections in both atlases clearly show this grouping based on mean C1R/S expression (Human, Figure S13c, d; Mouse, Figure S14c, d). In analysis of gene expression in heatmaps of the human atlases across cell-types and tissues C1R, C1S, and CFD expression was largely restricted to fibroblastic cells across tissue types, although we note expression of this gene seen in adipocytes in adipose tissue and marrow, a level of these proteins are also present in PVL's and SMC's across tissue, and liver epithelial expression (Figure S13e). In the mouse atlas we find C1R and C1S are widely expressed across tissue-types in fibroblasts, with trace expression on PVL's / SMC's, (Figure S14e), however no adipocyte or liver epithelial expression is seen. In mice CFD expression is reduced/absent, indicating that while this cell-type is likely conserved, CFD expression is dominant in the human variant of the cell-type. Given the lack of expression of CFD in mouse tissue generally vs. C1R/S inclusion of CFD in the signature is unlikely to alter results in mice, and will improve labeling in humans. Overall, we thus find that C1R, C1S, and CFD mark a conserved fibroblast type between mice and humans.

##### *Three patterns of complement expression are seen qualitatively*

Alongside widespread peripheral expression of the C1Q complement genes in myeloid cell-types and C1R/S in fibroblast cells, we also note both CFD and C3 emerged as defining markers of fibroblast cell-types in either the mouse (C3), or human (CFD) atlas (Table S5). We thus sought to understand expression of the full-set of complement family genes in scREF and scREF-mu, across the standardized cell-types to determine which cell-types express complement genes generally. Analysis of the cross-aligned atlas indicated quantitatively that three groupings emerged, alongside outlier behavior in C4A, C4B, CFI and CFP expression (main text). To further validate this finding we plotted complement family expression grouped by cell-type and tissue-type, averaged across the datasets in scREF (Figure S13e), and scREF-mu (Figure S14e). Here we confirmed that the hepatocyte expressed group (C5, C6, C8 components, C9 and CFB) is highly restrictive to hepatocytes in both species, though we note C6 can also be expressed in other liver cell-types, and CFB shows expression in non-liver epithelial cell-types. We also confirm that C2 is predominantly myeloid expressed and C7, C3, CFD, and CFH are principally fibroblast expressed, however a level of divergence between expression levels in mice and humans is seen for these proteins. Finally we assessed expression of outlier complement expression patterns and noted Placental/Uterine C4A/B expression, dendritic/ monocyte selective CFP expression, and divergent CFI expression, with endothelial cells expressing this gene in humans, while liver hepatocytes uniquely express this gene in mice (Figure 4F, Figure S13e, S14e). Overall this analysis thus further supports the existence of abundant complement expressing macrophage and fibroblast types in both humans and mice.

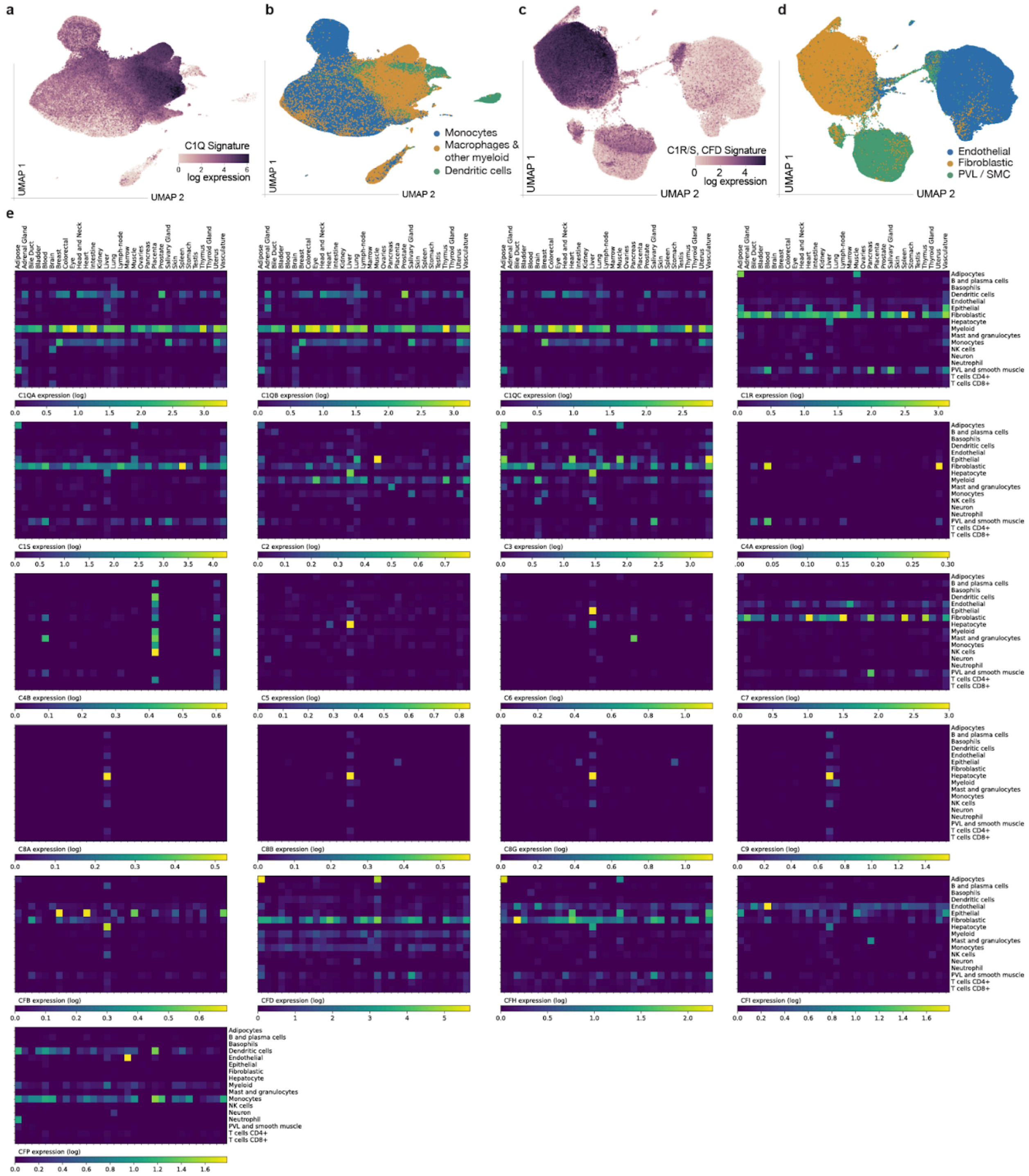

**Figure S13. Expression of complement family members across the full set of standardized cell-types and tissues in the human body, calculated from scREF:** a) UMAP projection of human myeloid cell-types coloured by C1Q signature score, calculated as the mean of the Log (read count +1) for C1Q genes C1QA, C1QB, and C1QC; b) The same projection as (a) but coloured by cell-type lineages; c) UMAP projection of human fibroblast, endothelial, and PVL/SMC cell-types coloured by C1R/S signature score, calculated as the mean of the Log (read count +1) for C1R, C1S; d) The same projection as (b) but coloured for cell-type label; e) Heatmaps of the log(read-count + 1) values of secreted complement pathway members, averaged for each standardized author cell-type and specific tissue-type for the human scREF atlas.

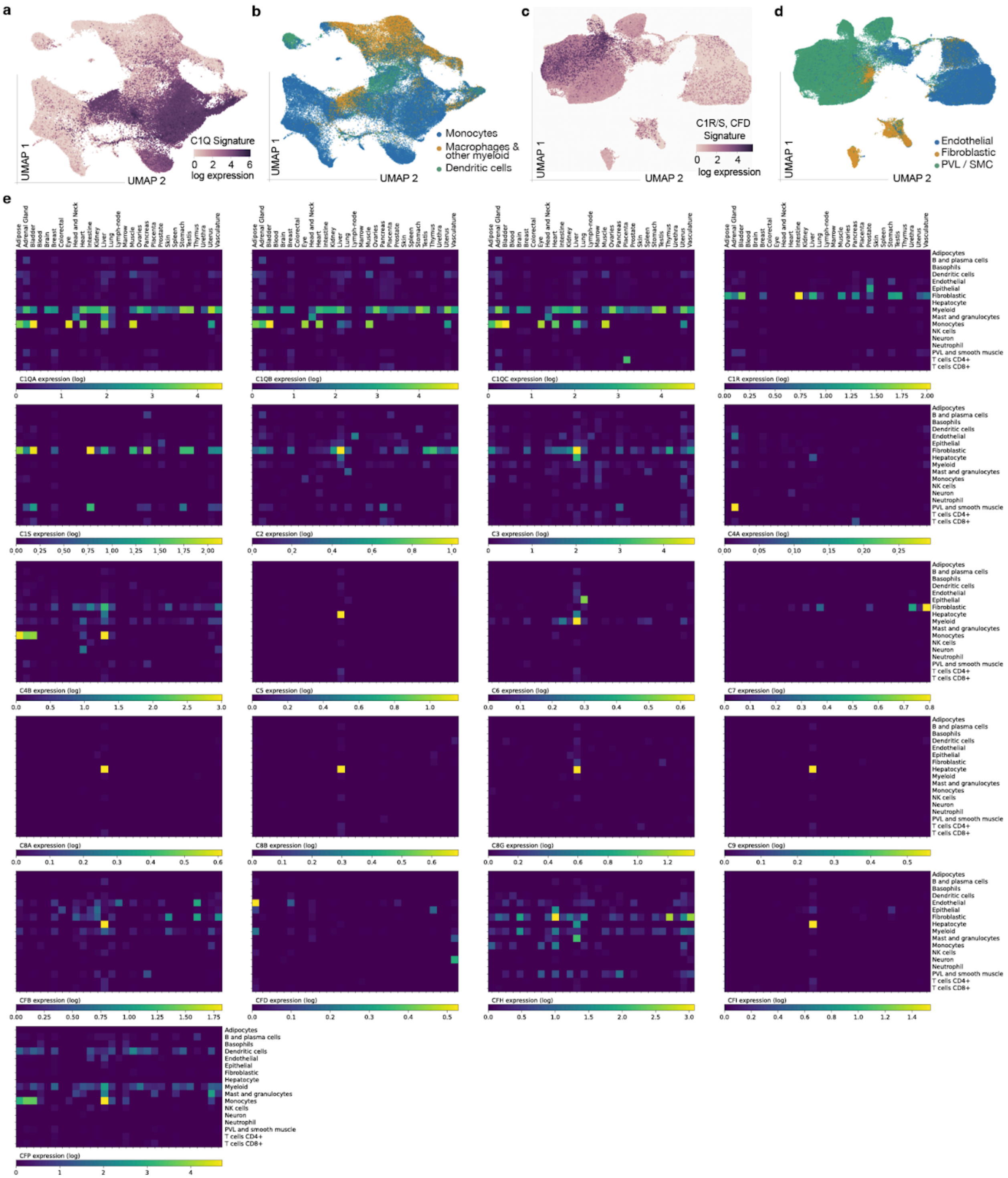

**Figure S14. Complement expressing macrophage and fibroblast types are present across mouse tissues:** a) UMAP projection of murine myeloid cell-types coloured by C1Q signature score, calculated as the mean of the Log (read count +1) for C1Q genes C1QA, C1QB, and C1QC; b) The same projection as (a) but coloured by cell-type lineages; c) UMAP projection of murine fibroblast, endothelial, and PVL/SMC cell-types coloured by C1R/S signature score, calculated as the mean of the Log (read count +1) for C1R, C1S; d) The same projection as (b) but coloured for cell-type label; e) Heatmaps of the log(read-count + 1) values of secreted complement pathway members, averaged for each standardized author cell-type and specific tissue-type for the murine scREF-mu atlas.

##### *Trained BA-ScVI can align a human atlas without further training or fine-tuning*

The high quality of alignment seen following the training of BA-scVI on scMARK and scREF provided conviction that BA-scVI could be valuable for *de novo* alignment of new human scRNA data, without further training, at scale. Specifically, where scVI requires the injection of batch ID vectors into both the encoder and decoder layers, preventing the alignment of new data without additional training (Lopez et al. 2018). BA-scVI, in contrast, only uses batch ID vectors to train the discriminator at train time (Figure 1b, Figure S5b). In line with this early tests demonstrated that BA-scVI trained on scMARK could be used to align new (never seen) data. Specifically, we predicted cell-types on a 12th dataset (Wu et al. 2021), held out at train time using BA-scVI trained on the 11 datasets present in scMARK. This provided initial confidence in this approach since UMAP projections show the accurate alignment of this previously unseen dataset (Figure S15a).

To test how well a trained BA-scVI model can perform de novo embedding at scale and across a wide variety of tissue-types we trained BA-scVI from scratch on scREF, but lacking the Tabula Sapiens atlas (scREF-TS) (Tabula Sapiens Consortium* et al. 2022), which comprises ~500k cells from 24 tissues across the human body (~220k cells in the downsampled training atlas). Notably this dataset has 171 unique original author labels defined, and 40 unique labels after standardization (more than any other study). This study also forms 18% of our stratified benchmark, as it has such a diverse array of tissues. Following *de novo* alignment we qualitatively saw a high degree of overlap also using a UMAP model trained on scREF-TS (Figure S15b), as well as clear grouping of both our standardized labels and author labels (Figure S15c, d). Quantitatively we first tested how well our embedding space captured the space used to label the cells in the original study. We used a K-nearest neighbor model trained on one half of the dataset to predict author labels on the other half of the dataset (Methods), here we obtained a 71% accuracy rate in predicting the original 171 author labels, indicating a high degree of preservation in cell-type label groupings between our embeddings and the original author embeddings used for cell-type labeling. This score is on par with studies that optimize models specifically for this task e.g., (Ghaddar and De 2022). With our standardized labels we achieve an even higher accuracy of 86%. Next we looked at our ability to predict standardized labels in Tabula Sapiens using scREF-TS, here we achieved an accuracy of 69%, notably less than the internal score, but still high given the 40 unique labels in this study. Finally we looked at the KNI and RbNI scores for this study, here we found that in the original scREF alignment we obtained a KNI and RbNI score of 0.648 and 0.619. In contrast on scREF-TS the KNI and RbNI scores decreased to 0.430 and 0.502 respectively. This indicates that while the trained BA-scVI model is accurate for *de novo* cell-type labeling and embedding, care should be taken to prior to performing cross dataset analysis of *de novo* aligned results, and inclusion of datasets at train time is preferred- we thus leave this capability out of the online tool we provide. We consider that with additional large datasets such as the Tabula Sapiens, yet better *de novo* KNI and RBNI scores will likely be obtained, and a consensus can be reached enabling *de novo* cross dataset comparison.

##### *BA-scVI enables the alignment of in vitro data to human tissue data*

To further confirm that BA-scVI is accurately aligning cell-types, and that author cell-type labels can act as a valuable ground-truth, we sought to align cell-type with a known ground truth. To do this we generated an *in vitro* scRNA dataset from cancer epithelial cells, normal fibroblasts, and cancer-associated fibroblasts (Methods) and performed alignment to the scMARK benchmark, since unlike the scREF benchmark it contains cancer epithelial cells, normal fibroblasts and cancer-associated fibroblasts. Qualitatively in UMAP projections we found that cancer epithelial and fibroblast cell-types aligned well to scMARK cancer cell and fibroblast clusters, although distinct sub-clusters were formed by tissue and in vitro cell-types (Figure S15e). Quantitatively, (99.2%) of in vitro cancer cells were correctly predicted while (95.5%) of fibroblasts were correctly predicted; here a KNN was used due to the presence of large, biological variation between the in-vitro cell-culture scRNA data and tissue cell data in scMARK (Methods). Overall this supports the notion that BA-scVI is accurate at aligning and predicting the label of scRNA data of known ground-truth, and also further supports the notion that standardized author labels are an accurate round truth. Of note, by not using the *in vitro* scRNA data at train time these insights are not biased by the learning of an alignment between in vitro and tissue data sources, and thus could be used to optimize model relevance.

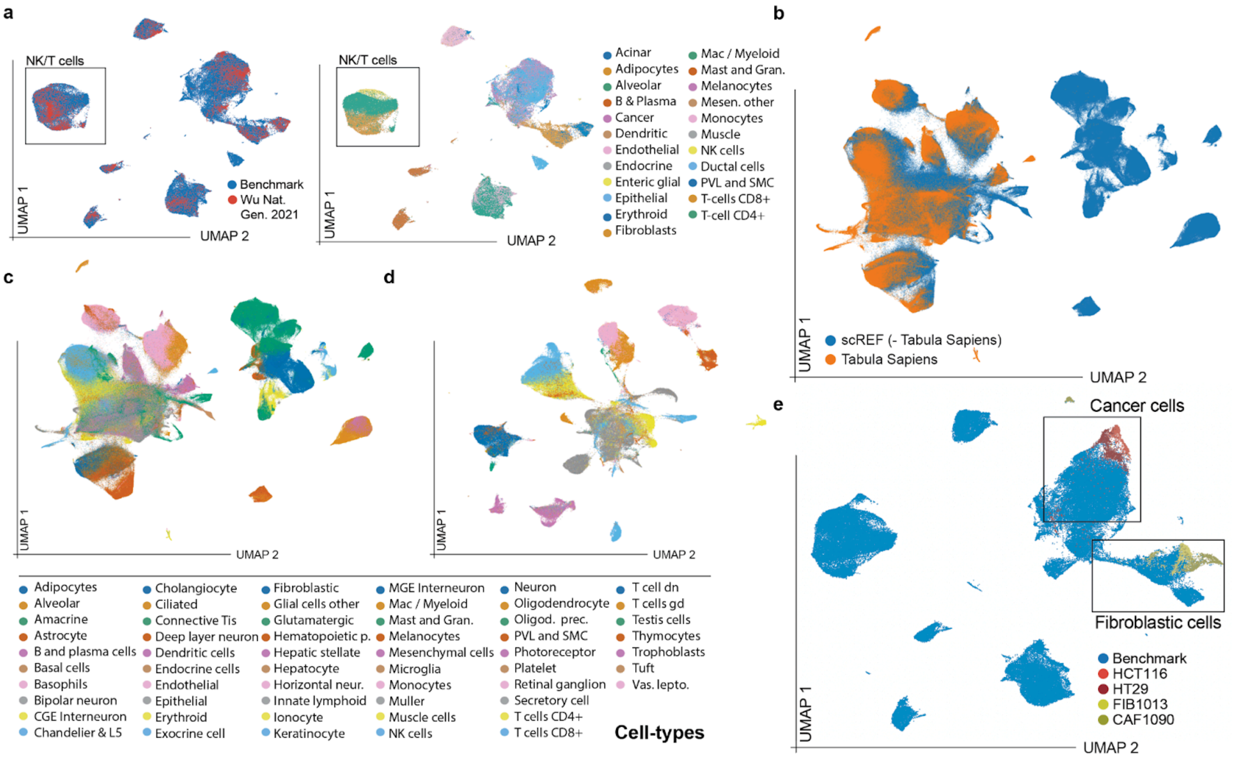

**Figure S15. Ba-scVI enables *de novo* alignment of new scRNA data:** a) UMAP projection of BA-scVI trained on scMARK and used to align a fully held out dataset (Wu, 2021) to scMARK, the overlap between dataset is shown in the left panels, cell-type labels in the right - highlighting e.g., effective overlap and separation of NK and T-cell types / subtypes; b) UMAP projection of the Tabula Sapiens atlas (orange) and remaining datasets in scREF (blue) where *de novo* alignment has been performed by BA-scVI following training on the remaining scREF datasets; c) The same UMAP projection as (b) coloured by standardized author label; d) A UMAP projection of the Tabula Sapiens atlas only (using *de novo* embeddings), coloured by standardized cell-type; e) UMAP projection of *in vitro* data aligned with BA-scVI *de novo* to the scMARK dataset, aligned data for four cell-lines is shown, namely HT29, HCT116, FIB1013, and CAF1090.

#### *Supplemental Methods (in vitro data generation)*

***in vitro cell culture:*** Human colon cancer cell lines HT-29 and HCT 116 were purchased from ATCC (Rockville, USA) and primary human normal (FIB1013) and cancer-associated fibroblasts (CAF1090) were purchased from BioIVT (Westbury, NY). All cells were kept at low passage (p1-5) and maintained in MEM medium (Hyclone cat#SH30244.01) supplemented with 10% FBS (ThermoFisher Cat#A5209402) , 2mM Glutamax (ThermoFisher Cat#35050079) , 1% Non-essential amino acids (Thermo Fisher Cat# 11140050), 1mM sodium pyruvate (Hyclone cat# ) SH30239.01 and 1% pen-strep (Hyclone cat#SV30010). Co-culture conditions including seeding density were optimized to yield an approximately 50:50 ratio of cancer to fibroblast cells after 48 hours of co-culture on gelatin coated plates. The following cell numbers were seeded onto gelatin-coated plates either as monocultures or mixed together as co-cultures: HT-29: 2.25e5 cells, HCT116: 1.25e5 cells, CAF1090: 6.0e5 cells, FIB1013: 1.0e6 cells. Cultures were established overnight and then switched to complete media with 0.5% FBS. After 48 hours, preparation of cell suspensions for single-cell RNA expression analysis were prepared by disassociation with Accutase™ Cell Dissociation Reagent (Stemcell Technologies Inc. cat# 07920).

***in vitro sequencing:*** Cell suspensions from each mono or co-culture were labeled with 3' CellPlex Multiplexing Solution (10x Genomics). Single-cell RNA-seq (scRNA-seq) was then performed at the Toronto UHN Princess Margaret Genomics Centre (PMGC) for droplet-based microfluidics using the 10x Chromium Single Cell platform (Single Cell 3′ v3, 10x Genomics, USA). Droplets of the cellular suspensions, reverse transcription master mix, and partitioning oil were mixed, loaded onto a single cell chip, and processed on the Chromium Controller. Reverse transcription was performed, and cDNA was amplified using a BioRad C1000 Touch thermal-cycler, with cDNA size selected using SpriSelect beads (Beckman Coulter, USA). An Agilent Bioanalyzer High Sensitivity DNA chip was used to analyze cDNA for qualitative control purposes; cDNA was then fragmented using the proprietary fragmentation enzyme blend for 5 min at 32°C, followed by end repair and A-tailing at 65°C for 30 min. DNA was double-sided size selected using SpriSelect beats. Sequencing adaptors were ligated to the cDNA at 20°C for 15 min cDNA was amplified using a sample-specific index oligo as primer, followed by another round of double-sided size selection using SpriSelect beads. Final libraries were analyzed on an Agilent Bioanalyzer High Sensitivity DNA chip for qualitative control purposes. Libraries were sequenced on a HiSeq 4000 Illumina platform targeting 50,000 reads per cell.

***in vitro single cell analysis:*** Base calls were converted to reads using the Cell Ranger (10X Genomics; version 3.1), aligned against either the GRCh38 v3.0.0 human reference genome.Cell Ranger's count function was set to SC3Pv3 chemistry and 5,000 expected cells per sample. Hashtag oligos (HTOs) were demultiplexed using Seurat's implementation HTODemux. Briefly, k-medoid clustering is performed on the normalized HTO values, after which a 'negative' HTO distribution is calculated. For each HTO, the cluster with the lowest average value is treated as the negative group, and a negative binomial distribution fits this cluster. Using the 99% quantile of this distribution as a threshold, each cell is classified as positive or negative for each HTO.

Cell barcodes representative of quality cells were differentiated from apoptotic cell barcodes or background RNA based on a threshold of having at least 200 unique transcripts profiled less than 100,000 total transcripts and less than 10% of their transcriptome of mitochondrial origin. Unique molecular identifiers (UMIs) from each cell barcode were retained for all downstream analyses, normalized with a scale factor of 10,000 UMIs per cell, and subsequently log(X+1) transformed using the R package Seurat (version 3.1.1; (Butler et al. 2018)). The first 15 principal components of the aggregated data were then used for uniform manifold approximation and projection (UMAP) analysis.

***KNN evaluation of in vitro data:*** *in vitro* cell-types were classified using a K-nearest neighbor model as per scikit-learn (Pedregosa et al. 2012), here k=25 neighbors were used, alongside default parameters. The model was trained to the full set of labels as defined in scMARK.

#### *Supplemental References*

Azizi, Elham, Ambrose J. Carr, George Plitas, Andrew E. Cornish, Catherine Konopacki, Sandhya Prabhakaran, Juozas Nainys, et al. 2018. "Single-Cell Map of Diverse Immune Phenotypes in the Breast Tumor Microenvironment." *Cell* 174 (5): 1293–1308.e36.

Butler, Andrew, Paul Hoffman, Peter Smibert, Efthymia Papalexi, and Rahul Satija. 2018. "Integrating Single-Cell Transcriptomic Data across Different Conditions, Technologies, and Species." *Nature Biotechnology* 36 (5): 411–20.

Büttner, Maren, Zhichao Miao, F. Alexander Wolf, Sarah A. Teichmann, and Fabian J. Theis. 2019. "A Test Metric for Assessing Single-Cell RNA-Seq Batch Correction." *Nature Methods* 16 (1): 43–49.

Cui, Haotian, Chloe Wang, Hassaan Maan, Kuan Pang, Fengning Luo, Nan Duan, and Bo Wang. 2024. "scGPT: Toward Building a Foundation Model for Single-Cell Multi-Omics Using Generative AI." *Nature Methods*, February. https://doi.org/10.1038/s41592-024-02201-0.

Dzionek, Andrzej, Yoshiaki Sohma, Jun Nagafune, Marina Cella, Marco Colonna, Fabio Facchetti, Gritt Günther, et al. 2001. "BDCA-2, a Novel Plasmacytoid Dendritic Cell–specific Type II C-Type Lectin, Mediates Antigen Capture and Is a Potent Inhibitor of Interferon α/β Induction." *The Journal of Experimental Medicine* 194 (12): 1823–34.

Furlan, Alessandro, and Igor Adameyko. 2018. "Schwann Cell Precursor: A Neural Crest Cell in Disguise?" *Developmental Biology* 444 Suppl 1 (December):S25–35.

Gabitto, Mariano I., Kyle J. Travaglini, Victoria M. Rachleff, Eitan S. Kaplan, Brian Long, Jeanelle Ariza, Yi Ding, et al. 2023. "Integrated Multimodal Cell Atlas of Alzheimer's Disease." *Research Square*, May. https://doi.org/10.21203/rs.3.rs-2921860/v1.

Ghaddar, Bassel, and Subhajyoti De. 2022. "Census: Accurate, Automated, Deep, Fast, and Hierarchical scRNA-Seq Cell-Type Annotation." *bioRxiv*. https://doi.org/10.1101/2022.10.19.512926.

Hao, Yuhan, Stephanie Hao, Erica Andersen-Nissen, William M. Mauck 3rd, Shiwei Zheng, Andrew Butler, Maddie J. Lee, et al. 2021. "Integrated Analysis of Multimodal Single-Cell Data." *Cell* 184 (13): 3573–87.e29.

Hie, Brian, Bryan Bryson, and Bonnie Berger. 2019. "Efficient Integration of Heterogeneous Single-Cell Transcriptomes Using Scanorama." *Nature Biotechnology* 37 (6): 685–91.

Karlsson, Max, Cheng Zhang, Loren Méar, Wen Zhong, Andreas Digre, Borbala Katona, Evelina Sjöstedt, et al. 2021. "A Single-Cell Type Transcriptomics Map of Human Tissues." *Science Advances* 7 (31). https://doi.org/10.1126/sciadv.abh2169.

Kasprzak, Aldona. 2021. "Somatostatin and Its Receptor System in Colorectal Cancer." *Biomedicines* 9 (11). https://doi.org/10.3390/biomedicines9111743.

Kingma, Diederik P., and Max Welling. 2013. "Auto-Encoding Variational Bayes." *arXiv [stat.ML]*. arXiv. http://arxiv.org/abs/1312.6114v11.

Kiselev, Vladimir Yu, Andrew Yiu, and Martin Hemberg. 2018. "Scmap: Projection of Single-Cell RNA-Seq Data across Data Sets." *Nature Methods* 15 (5): 359–62.

Korsunsky, Ilya, Nghia Millard, Jean Fan, Kamil Slowikowski, Fan Zhang, Kevin Wei, Yuriy Baglaenko, Michael Brenner, Po-Ru Loh, and Soumya Raychaudhuri. 2019. "Fast, Sensitive and Accurate Integration of Single-Cell Data with Harmony." *Nature Methods* 16 (12): 1289–96.

Kozareva, Velina, Caroline Martin, Tomas Osorno, Stephanie Rudolph, Chong Guo, Charles Vanderburg, Naeem Nadaf, Aviv Regev, Wade G. Regehr, and Evan Macosko. 2022. "Author Correction: A Transcriptomic Atlas of Mouse Cerebellar Cortex Comprehensively Defines Cell Types." *Nature* 602 (7896): E21.

Lopez, Romain, Jeffrey Regier, Michael B. Cole, Michael I. Jordan, and Nir Yosef. 2018. "Deep Generative Modeling for Single-Cell Transcriptomics." *Nature Methods* 15 (12): 1053–58.

McInnes, Leland, John Healy, and James Melville. 2018. "UMAP: Uniform Manifold Approximation and Projection for Dimension Reduction." *arXiv [stat.ML]*. arXiv. http://arxiv.org/abs/1802.03426.

Murray, Liisa, Yang Xi, and John W. Upham. 2019. "CLEC4C Gene Expression Can Be Used to Quantify Circulating Plasmacytoid Dendritic Cells." *Journal of Immunological Methods* 464 (January):126–30.

Nagatake, Takahiro, Harumi Fujita, Nagahiro Minato, and Yoko Hamazaki. 2014. “Enteroendocrine Cells Are Specifically Marked by Cell Surface Expression of Claudin-4 in Mouse Small Intestine.” *PloS One* 9 (6): e90638.

Pedregosa, Fabian, Gaël Varoquaux, Alexandre Gramfort, Vincent Michel, Bertrand Thirion, Olivier Grisel, Mathieu Blondel, et al. 2011. “Scikit-Learn: Machine Learning in Python.” *The Journal of Machine Learning Research* 12:2825–30.

———. 2012. “Scikit-Learn: Machine Learning in Python.” *arXiv [cs.LG]*. arXiv. https://www.jmlr.org/papers/volume12/pedregosa11a/pedregosa11a.pdf?ref=https:/.

Rousseeuw, Peter J. 1987. “Silhouettes: A Graphical Aid to the Interpretation and Validation of Cluster Analysis.” *Journal of Computational and Applied Mathematics* 20 (November):53–65.

Shigematsu, Hirokazu, Boris Reizis, Hiromi Iwasaki, Shin-Ichi Mizuno, Dan Hu, David Traver, Philip Leder, Nobuo Sakaguchi, and Koichi Akashi. 2004. “Plasmacytoid Dendritic Cells Activate Lymphoid-Specific Genetic Programs Irrespective of Their Cellular Origin.” *Immunity* 21 (1): 43–53.

Siletti, Kimberly, Rebecca Hodge, Alejandro Mossi Albiach, Ka Wai Lee, Song-Lin Ding, Lijuan Hu, Peter Lönnerberg, et al. 2023. “Transcriptomic Diversity of Cell Types across the Adult Human Brain.” *Science* 382 (6667): eadd7046.

Tabula Sapiens Consortium*, Robert C. Jones, Jim Karkanias, Mark A. Krasnow, Angela Oliveira Pisco, Stephen R. Quake, Julia Salzman, et al. 2022. “The Tabula Sapiens: A Multiple-Organ, Single-Cell Transcriptomic Atlas of Humans.” *Science* 376 (6594): eabl4896.

Theodoris, Christina V., Ling Xiao, Anant Chopra, Mark D. Chaffin, Zeina R. Al Sayed, Matthew C. Hill, Helene Mantineo, et al. 2023. “Transfer Learning Enables Predictions in Network Biology.” *Nature* 618 (7965): 616–24.
